# The P2X7 receptor is an intermediate in the retinoic acid-signaling pathway that induces neuronal remodeling in retinitis pigmentosa

**DOI:** 10.64898/2026.05.27.728267

**Authors:** Michael Telias, Leonor Afrima, Jared Endres, Bristol Denlinger, Richard H. Kramer

## Abstract

Photoreceptor degeneration drives electrophysiological remodeling in downstream retinal neurons, including hyperactive firing of retinal ganglion cells (RGCs) that degrades residual vision. Although Retinoic Acid (RA) and its receptor (RAR) are known to upregulate ion channel genes associated with RGC hyperactivity, these genes lack canonical RAR binding sites required for direct transcriptional regulation. Here, we identify P2X7 receptor (P2X7R) as a key intermediary in RA-induced remodeling in the rd1 mouse model of retinitis pigmentosa. Genetic deletion of P2X7R prevents the upregulation of RA-responsive genes and abolishes RGC hyperactivity. Loss of P2X7R also eliminates RGC hyperpermeability, as measured by uptake of an otherwise membrane-impermeant dye. Acute pharmacological inhibition of P2X7R suppresses hyperpermeability without affecting hyperactivity, supporting an indirect signaling mechanism rather than a direct electrophysiological role. Notably, elevation of resting Ca²⁺ is absent in P2X7R-deficient cells, implicating Ca²⁺-dependent gene expression as a link between P2X7R-mediated hyperpermeability and RGC hyperactivity. Together, these findings establish P2X7R as a critical intermediate in retinal remodeling and a potential therapeutic target for preserving vision in retinitis pigmentosa.

**Graphical Abstract:** 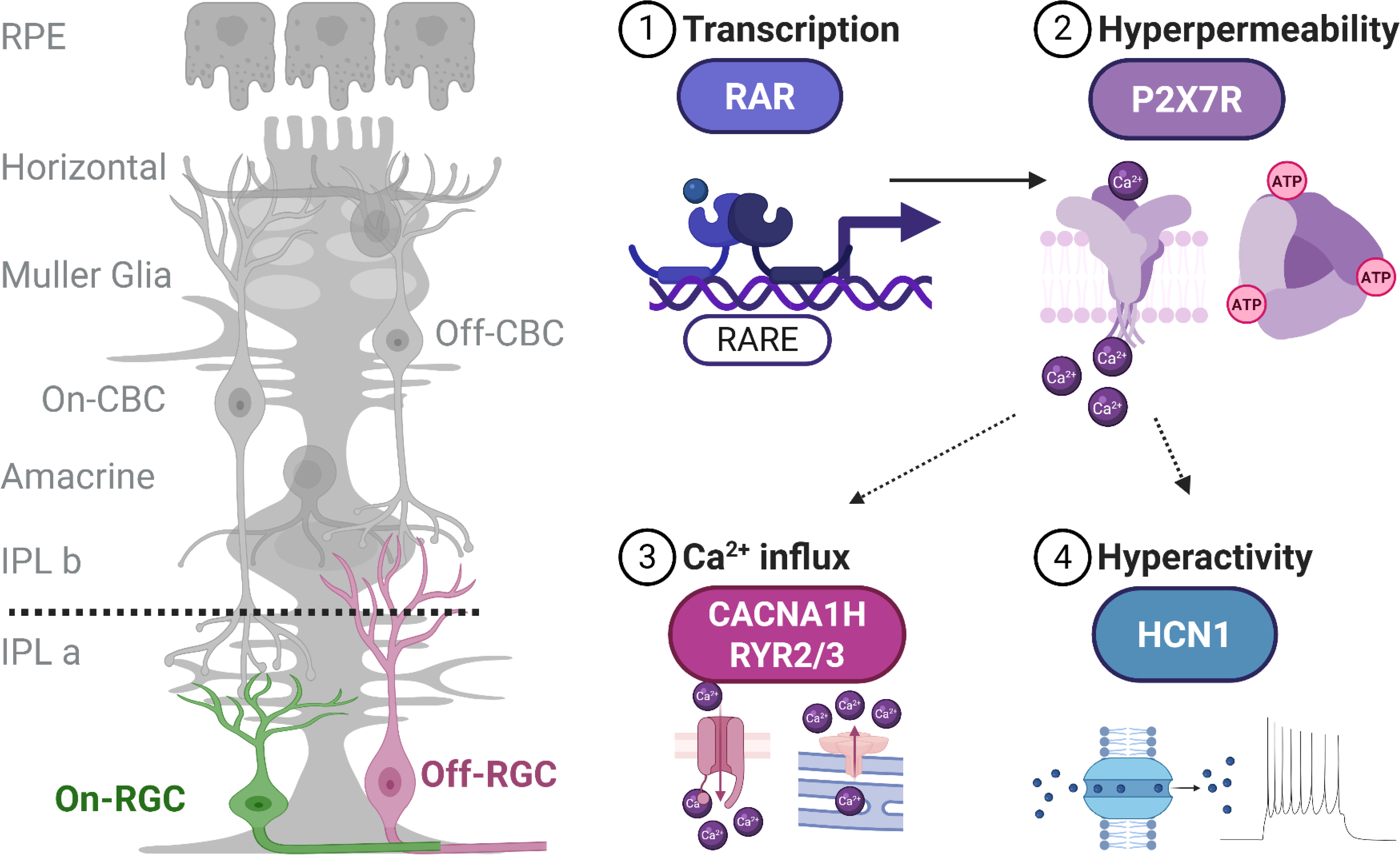

## Introduction

Retinal photoreceptors degenerate in inherited retinal disorders such as retinitis pigmentosa (RP), and non-inherited disorders, such as age-related macular degeneration (AMD), leading to progressive vision impairment and blindness. The loss of light-elicited signals from rods and cones is the primary reason for vision loss in AMD and RP, but changes in the physiology and morphology of downstream retinal neurons corrupt processing of visual information, also contributing to visual decline [1, 2]. The neural network of the retina has a remarkable ability to compensate for partial loss of photoreceptors [3–5], but the massive degeneration characteristic of RP and AMD also has pathophysiological consequences on downstream neurons, interfering with normal circuit function. ON-type retinal bipolar cells (BCs) lose over 75% of their voltage-gated calcium conductance, decreasing the amount of neurotransmitter released onto postsynaptic neurons and altering the strength and dynamics of synaptic transmission [6]. Some types of RGCs react to photoreceptor loss by spontaneously firing action potentials at a high rate, diminishing their ability to encode light responses initiated by surviving rod and cone photoreceptors [7]. These effects propagate through the visual system to the primary visual cortex, where corruption of input signals interferes with feature detection by pyramidal neurons, reducing orientation-selectivity and decreasing the reliability of their responses to complex features in naturalistic visual scenes [2]. The net consequence of these physiological changes is reduced perceptual ability, revealed with a behavioral visual recognition task carried out on the rd10 slowly degenerating mouse model of RP.

The signaling molecule that triggers all these aspects of remodeling is retinoic acid (RA), a vitamin A metabolite that regulates gene transcription (*canonical pathway*) [8, 9] and translation (*non-canonical pathway*) [10–12]. Several lines of evidence implicate RA [7]. Artificially introducing RA into the eye mimics aspects of retinal remodeling in wild-type mice. Inhibiting RA synthesis by the enzyme RALDH or RA signaling by the nuclear RA receptor (RAR) suppresses retinal remodeling in rd1 and rd10 mice. Finally, elevated RA-induced gene expression has been detected in RGCs from rd1 mice with a fluorescent protein reporter gene [7]. These experiments establish that RA is necessary and sufficient for pathophysiological remodeling of RGCs, but exactly how RA-induced gene expression leads to hyperactive RGC firing has remained unclear.

One surprising feature of remodeling is an increase in the permeability of the RGC cell membrane to organic cations that are normally impermeant [13]. The cationic fluorescent dye Yo-Pro-1 is excluded from RGCs in healthy wild-type retinas, but the dye permeates into RGCs in rd1 and rd10 retinas, brightly staining their nuclei upon binding to DNA. Azobenzene photoswitches, such as BENAQ, which contain a positively charged quaternary ammonium group, are also excluded from RGCs in wild-type retinas, but they also permeate RGCs in photoreceptor-degenerated retinas. Once inside the cell, *trans* BENAQ blocks the pore of voltage-gated ion channels, suppressing firing, but light photoisomerizes the molecule to the cis form, unblocking the pore. In this manner, BENAQ and other photoswitches confer light-sensitive control over action potential firing in RGCs. This property has made BENAQ a promising drug candidate for restoring vision in RP patients with RP, with clinical trials evaluating safety and efficacy underway in Australia (KIO-301, Kiora Pharmaceuticals).

The membrane conduit that allows Yo-Pro-1 and BENAQ to enter cells is the P2X receptor, a ligand-gated ion channel that is activated by extracellular ATP. P2X receptors (*p2rx*) are non-selective cation channels, with the open state conducting small physiological cations including Na^+^, K^+^, and Ca^+2^. However, some P2X receptor subtypes, including *p2rx7*, have a “macropore” that is sufficiently large to accommodate larger organic cations, including Yo-Pro-1 and BENAQ [13]. The structural basis and functional significance of the macropore remain unclear, but these receptors have been implicated in activating downstream signaling cascades, including the nuclear factor of activated T-cells (NFAT) family of transcription factors [14], and mitogen-activated protein kinase (MAPK) [15]. However, the mechanistic contribution of *p2rx7* to inner retinal remodeling and RGC hyperactivity has not been addressed.

Here we show that expression of *p2rx7* and functional upregulation of P2X7 receptors (P2X7Rs) in the inner retina is necessary and sufficient to induce membrane hyperpermeability and spontaneous retinal hyperactivity. We further demonstrate that degeneration-dependent upregulation of *p2rx7* is correlated with the upregulation of dozens of genes related to excitability, calcium-dependent signaling and voltage-gated ion channels. Specifically, we found that in degenerated retinas, *p2rx7* is necessary for transcriptional and functional upregulation of hyperpolarization-activated cyclic nucleotide-gated channels (HCN1) exclusively in Off-RGCs; suggesting that inner retinal remodeling, triggered by RA/RAR and mediated by P2X7Rs, results in asymmetric functional corruption, affecting Off RGCs more than On-RGCs. Our study identifies *p2rx7* as a potential target in mitigation of visual loss in IRDs, and sheds light on the molecular and physiological mechanisms behind circuit corruption in neuronal maladaptive plasticity.

## Results

### Degeneration-induced upregulation of P2X7Rs is downstream to RA-signaling

Studies have shown that loss of rods and cones in mouse models of RP is correlated with increased transcription [13], translation [16], and activation [7, 13] of P2X7Rs in the surviving inner retina. Here, we used immunofluorescence to visualize the expression pattern of P2X7R in retinal slices from adult (P60-90) WT (**Fig. 1A**) and rd1 (**Fig. 1B**) mice. We quantified the fluorescence of anti-P2X7R immunolabeling (relative to unstained controls) across layer-specific regions of interest (ROIs, **Fig. 1A**, yellow boxes). We used DAPI (blue) to reveal all cell nuclei and co-labeling with the RGC-specific marker RBPMS (red) as an anatomical hallmark [17–19]. Overall, we measured an increase in P2X7R immunolabeling of the inner nuclear layer (INL, **Fig. 1C**) and the inner plexiform layer (IPL, **Fig. 1D**), and a non-significant increase in the ganglion cell layer (GCL, **Fig. 1E, p=0.07**). These results show that photoreceptor loss is correlated with increased *p2rx7* transcription and P2X7R translation in the inner retina.

**Figure 1.**
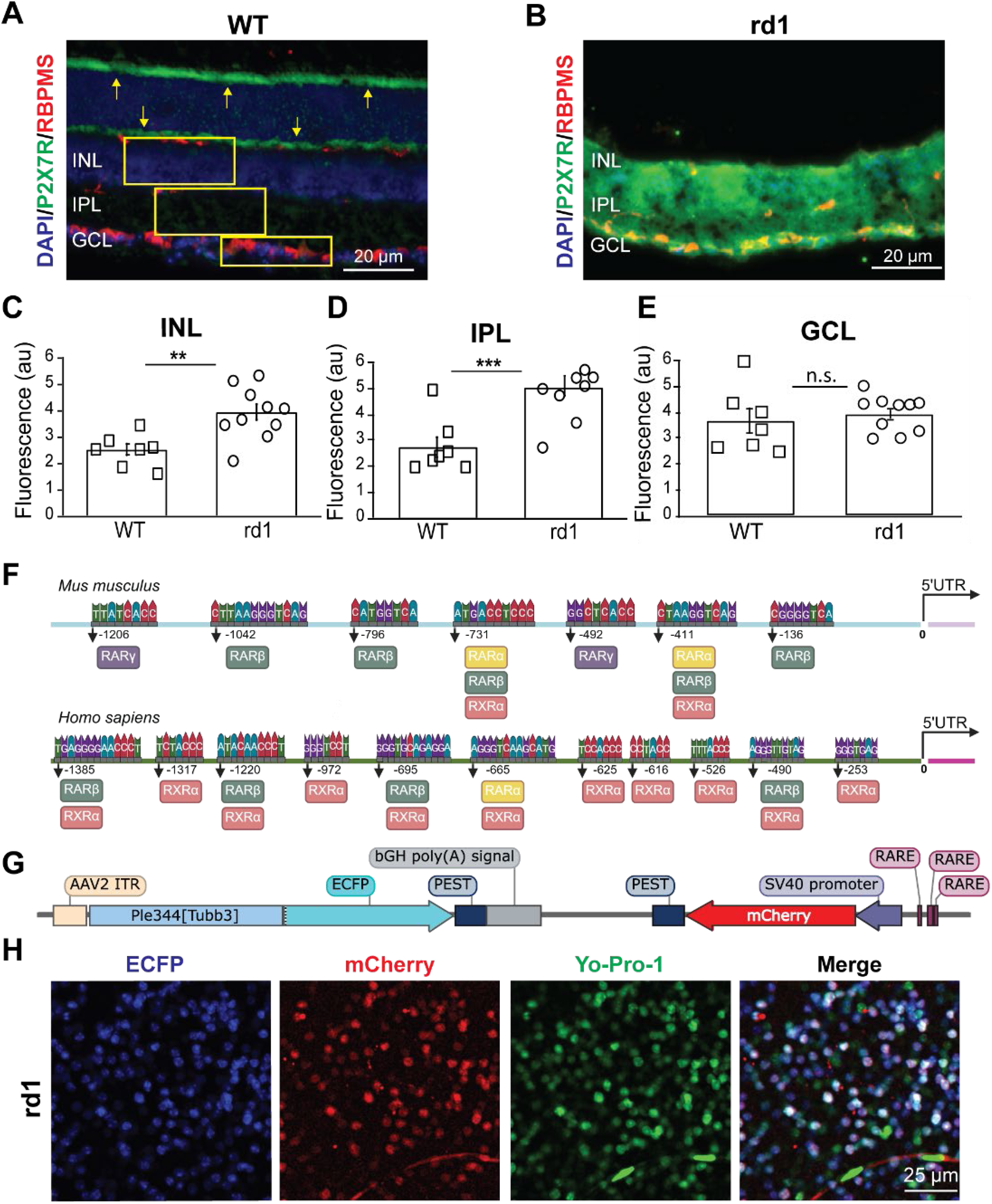
Degeneration- and RA-dependent upregulation of *p2rx7* in the rd1 inner retina. **A)** Immunofluorescence labeling of cell nuclei (DAPI, blue), P2X7R protein (green) and the retinal ganglion cell marker RBPMS (red) in a retinal slice from an adult WT mouse. Yellow arrows show two non-specific bands stained by the secondary antibody (see also **Fig. S1**) and yellow regions of interest illustrating the method employed to quantify fluorescence. INL – inner nuclear layer; IPL – inner plexiform layer; GCL – ganglion cell layer. **B)** Similar to (**A**), but in the retina of an adult rd1 mouse, fully degenerated. **C-E**) Quantification of P2X7R green fluorescence in all three regions of interest detailed above. Images here had their brightness and contrast artificially enhanced for graphic purposes, but analysis was conducted on raw images without any manipulation. Individual data points represent the average of 3-5 fields of view in a single retina from a single mouse (WT n=7 mice; rd1 n=10 mice, P60-90). Data are shown as mean ± SEM. **p<0.01, ***p<0.001, n.s. – non-significant; 1-tailed t-test after Shapiro-Wilk normality test. No outliners detected (Thompson Tau test). **F)** Illustration of the type, number and position of Retinoic Acid Response Element (RARE) sequences for all three RAR subtypes (α, β, γ), as well as for RXRα, in the first 1500 bp immediately upstream to the 5’ untranslated region of the P2RX7 promoter, depicting the mouse gene (*Mus musculus*) on the top, and the human homolog region on the bottom (*Homo sapiens*). **G)** Schematic sequence map for the RAR-reporter construct, including enhanced cyan fluorescent protein (ECFP) under the neuronal artificial mini-promoter Ple344 derived from Tubb3, the expression of red fluorescent protein (mCherry) under the regulation of the weak SV40 promoter and three repetitions of a RARE sequence. Fluorescent proteins express the degradation peptide PEST, to prevent overexpression. **H)** Images of a flat-mounted retinal sample, with the GCL facing up, dissected from a fully degenerated rd1 mouse after intravitreal delivery of the RAR-reporter and acute incubation with the P2X7R-dependent nuclear dye Yo-Pro-1.

To substantiate this conclusion, we analyzed the genomic sequence of the mouse and human *p2rx7* promoter and found many RAR-binding sites and retinoid orphan receptor (RXR)-binding sites within the first 1500 bp upstream from the transcription initiation codon [20, 21], suggesting that *p2rx7* transcription might be directly regulated by RAR/RXR (**Fig. 1F**). To confirm a direct link between RA-signaling, translational P2X7R upregulation, and functional receptor activation, we conducted two different experiments. First, we labeled *in-vivo* RGCs in rd1 mice, using a genetically encoded ratiometric fluorescent protein-based RAR-reporter encoding blue fluorescent protein to label infected cells and red fluorescent protein to reflect RAR-dependent transcription (**Fig. 1G**, intravitreal AAV2 delivery). When retinal samples were stained with the Yo-Pro-1 dye (**Fig. 1H**, green), over 90% of mCherry-positive cells in the GCL were Yo-Pro-1-positive, suggesting a strong correlation between high RAR-dependent transcription and *p2xr7* expression.

### Upregulation of P2X7R underlies RGC membrane hyperpermeability downstream to RA-signaling in degenerating retinas

To test the role of P2X7R in functional remodeling, we generated a double mutant mouse by crossbreeding the rd1 mouse (*Pde6β^rd1/rd1^*), with the *p2rx7* knock-out (ko) mouse (*P2rx7^-/-^*) (**Fig. 2A**). The genotype of the resulting mice, named ‘rd1-*p2rx7*ko’, shows biallelic loss of the *p2rx7* gene (**Fig. 2B**), and the expected loss of mRNA transcription (**Fig. 2C**) and protein translation (**Fig. 2G, green band in rd1-*p2rx7*ko**). As expected, RGC permeability to Yo-Pro-1 is completely absent in the rd1-*p2rx7*ko mice (**Fig. 2D,E**), further confirming the essential role of this receptor subunit in membrane hyperpermeability in the degenerated retina.

**Figure 2.**
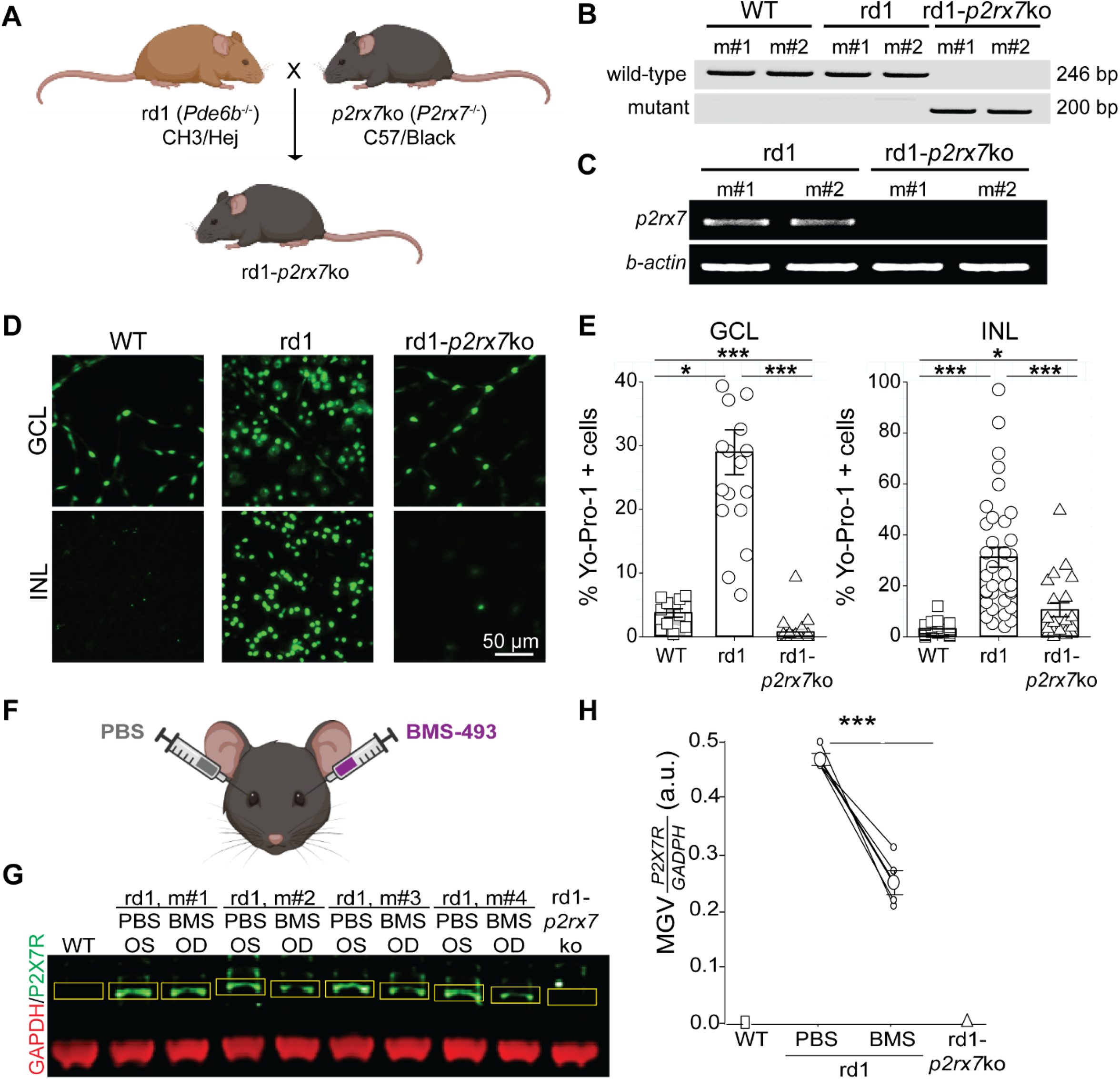
Genomic deletion of *p2rx7* in rd1 mice rescues membrane hyperpermeability. **A)** Illustration of breeding strategy to obtain double mutant rd1-*p2rx7*ko mice (for strain details see STAR Methods). **B)** Gel electrophoresis of PCR amplicons corresponding to the wild-type (top) and mutant (bottom) alleles of the mouse *p2rx7* gene in the original knock-out strain. Mice with no retinal degeneration (WT) were compared to rd1 and rd1-*p2rx7*ko (P21, n=2/condition). The single-nucleotide mutation in both Pde6b alleles in rd1 and rd1-*p2rx7*ko mice was confirmed by DNA sequencing. **C)** Gel electrophoresis of reverse transcription PCR amplicons for p2xr7 and b-actin demonstrate lack of *p2xr7* mRNA expression in rd1-*p2rx7*ko. RNA was purified from total retinal lysates in P60 mice (n=2/condition). **D)** Confocal images of flat-mounted retinas from WT, rd1 and rd1-p2rx7ko mice, incubated with Yo-Pro-1 dye, showing regions of interest corresponding to the ganglion cell layer (GCL, top) and the inner nuclear layer (INL). The GCL of WT and rd1-*p2rx7*ko retinas show Yo-Pro-positive pericytes in blood vessels. **E)** Quantification of the percentage of cells positively stained with Yo-Pro-1 out of all cells in each region of interest (DAPI, not shown). Data points correspond to a 200 µm x 200 µm area, 2 – 4 regions of interest/retina, 1-2 retinas/mouse, and 5-7 mice/strain, all at P60-90. Data are shown as mean ± SEM. *p<0.05, ***p<0.001, 2-tailed t-test after Shapiro-Wilk normality test. **F)** Illustration of intravitreal drug delivery of 2 µl solution containing 100 µM BMS-493 (BMS) in one eye, and contralateral vehicle control (PBS), repeated in 4 individual rd1 mice. **G-H)** Western blot gel bands for loading control GAPDH (red) and P2X7R (green) individual retinas from the left (OS) or right (OD) eye. Yellow boxes correspond to regions of interest used for quantitative analysis. Data are shown as mean ± SEM.***p<0.001; Mann-Whitney U test.

Next, we quantified P2X7R protein levels using Western blot (WB) in cell lysates obtained from single retinas, 3 days after intravitreal delivery of the potent pan-RAR antagonist BMS-493 (BMS) to one eye, and its vehicle (saline, PBS) control to the second eye (**Fig. 2F**). In rd1 retinas, WB analysis showed that a single intraocular injection of BMS-493 reduced P2X7R protein levels by half within 72 hrs. (**Fig. 2G,H**), as compared to the loading control protein (GAPDH) which showed equal quantities across samples (**Fig 2G**, **red bands**). In the same assay, the healthy retina of a WT mouse showed insignificant levels of P2X7R protein, similar to rd1-*p2rx7*ko, confirming previous observations of low to no expression of P2X7R in healthy retinal tissue [7, 13, 22]. We therefore conclude that photoreceptor loss unleashes RA-signaling within the inner retina, directly leading to the transcriptional, translational, and functional upregulation of P2X7R.

### P2X7Rs are necessary and sufficient to induce RGC spontaneous hyperactivity

Our published [7, 13, 22] and new data (**Fig 1,2**) all point to P2X7Rs being necessary and sufficient to induce RGC-membrane hyperpermeability, under direct regulation by RA-signaling during photoreceptor degeneration. This raises the possibility that P2X7Rs are also necessary and sufficient for spontaneous RGC-hyperactivity [2]. Multielectrode array (MEA) recordings of *ex-vivo* retinal samples show that genomic deletion of *p2rx7* rescued spontaneous RGC hyperactivity in the rd1 mouse (**Fig. 3A**). The average spontaneous firing rate of RGCs in rd1-*p2rx7*ko retinal samples was ∼50% of that for rd1 retinas (**Fig. 3B**). This was equally true whether the retina was bathed in saline, with or without a mixture of neurotransmitter receptor inhibitors (for glutamate-, GABA-, glycine-, and acetylcholine receptors) that isolates RGCs from chemical synaptic inputs (**Fig. 3C**). Hence RGC hyperactivity, induced by RA and made possible by P2X7R, can be attributed to changes in the intrinsic electrophysiological properties of RGCs. Importantly, while genomic deletion of *p2rx7* was sufficient to prevent hyperactivity, acute pharmacological blockade of P2X7R by the antagonist TNP-ATP was not (**Fig. 3B-C**), suggesting that sustained P2X7R expression and activation (in the range of hours to days) is needed to induce hyperactivity, rather than more immediate and transient mechanisms, such as cell depolarization (in the range of milliseconds to minutes).

**Figure 3.**
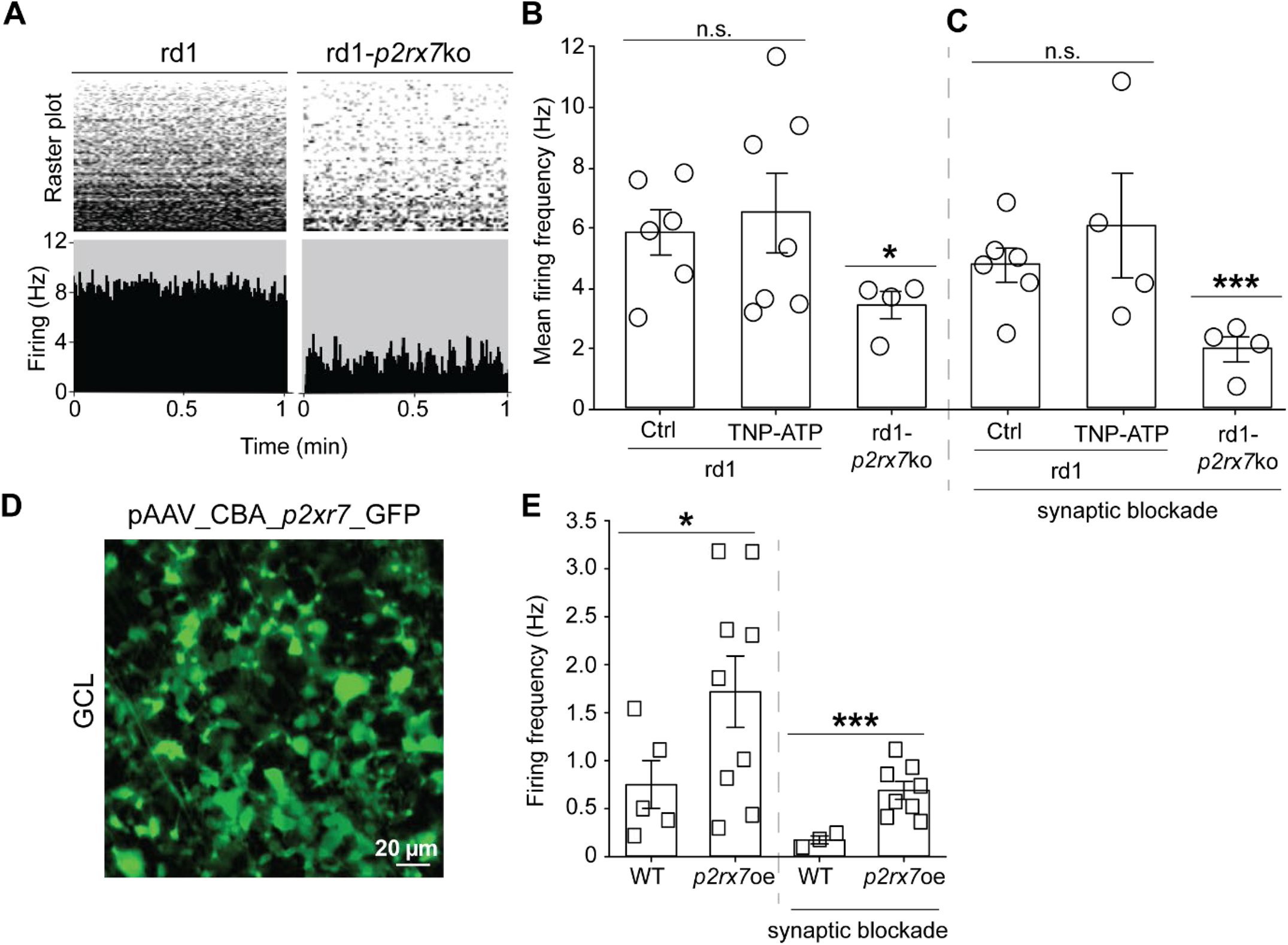
P2X7 receptors mediate degeneration-dependent RGC hyperactivity. **A)** Representative multielectrode array recording of spontaneous extracellular retinal activity in rd1 and rd1-*p2rx7*ko excised retinas, during 1 minute, under continuous perfusion with oxygenated saline at 34°C. Raster plots are shown on the top, and the combined firing activity (in Hz) of all detected units, shown on the bottom. **B)** Quantification of mean firing frequency for rd1 retinas in saline (Ctrl) or treated with a pan-P2X antagonist (TNP-ATP, 100 µM), and rd1-*p2rx7*ko retinas in saline. **C)** In a second experimental set, we repeated the same conditions while perfusing saline containing a cocktail of synaptic blockers (see STAR Methods). Data are shown as mean ± SEM. *p<0.05, ***p<0.001, n.s. – non-significant, 2-tailed t-test after Shapiro-Wilk normality test. **D)** Flat-mount fluorescent-light imaging of the GCL in a WT mouse infected intravitreally with an adeno-associated virus (AAV) delivering overexpression of *p2rx7* and green fluorescent protein (GFP) under the chicken beta actin (CBA) constitutive promoter. **E)** Quantification of spontaneous activity in the dark from the GCL of WT retinas treated with pAAV_CBA_P2RX7_GFP or not, under saline perfusion with or without synaptic blockers. Data are shown as mean ± SEM. *p<0.05, ***p<0.001, 2-tailed t-test after Shapiro-Wilk normality test.

To directly test whether *p2rx7* up-regulation is sufficient to induce RGC hyperactivity, we injected WT mice intravitreally with a viral construct encoding *p2xr7* under the strong chicken-beta actin promoter (CBA) (**Fig. 3D**). Overexpression of *p2rx7* (*p2rx7*oe) in WT retinas, without addition of exogenous ATP, was sufficient to induce a 2-3-fold-increase in the spontaneous firing rate of RGCs (**Fig. 3E**). Application of the neurotransmitter inhibitor cocktail, by itself, reduced the overall firing rate of RGCs, indicating that synaptic inputs are involved in setting the spontaneous firing rate. However, this input was insufficient to account for the effects of *p2rx7*oe on spontaneous activity (**Fig. 3E**, ‘synaptic blockade’). In summary, these results demonstrate that RA-induced *p2xr7* upregulation in the degenerated rd1 retina is necessary and sufficient to induce membrane hyperpermeability and RGC hyperactivity, and that these effects are brought about by downstream effectors in the RA-P2X7R signaling axis.

### *P2rx7* is necessary for degeneration-dependent changes in gene expression contributing to RGC hyperactivity

The P2x receptor antagonist TNP-ATP acts quickly (within 15 min) to block RGC hyperpermeability in rd1 retinas [7, 13], consistent with conduction of the dye through open P2X channels. Over the same time period, TNP-ATP had no effect on RGC hyperactivity (**Fig. 3B-C**). It is only when P2X7R is absent over a prolonged period, as in the rd1-*p2rx7*ko mouse, that hyperactivity is absent. This suggests that *p2rx7* plays a different role in hyperactivity. In this scenario, the P2X7R provides an intermediate signal, for example elevated intracellular Ca^2+^, that perhaps leads to altered expression of genes encoding voltage-dependent ion-channels responsible for hyperactivity, or perhaps proteins that directly or indirectly regulate those channels. To evaluate this possibility, we conducted an unbiased transcriptomic screen using next-generation sequencing of total retinal RNA from rd1-*p2rx7*ko and rd1 mice [23]. Principal component analysis and hierarchical clustering across rd1-*p2rx7*ko samples and the two rd1 sub-strains (C57 and C3H backgrounds) revealed distinct clustering of the double-mutant group, with substantial separation from both rd1 backgrounds, indicating that the observed transcriptional changes were driven by *p2rx7* deletion rather than genetic background (**Fig. S2A**,**B**). Transcriptome analysis identified 1,435 differentially expressed genes (ranked by p-value). DESeq2 analysis revealed that the majority of these genes were downregulated, while a smaller subset was upregulated (**Fig. 4A**,**B**; **Fig. S2C**). Gene ontology enrichment analysis identified 11 significantly upregulated pathways encompassing 76 genes, and 32 significantly downregulated pathways encompassing 612 genes. Prominent functional categories among the downregulated pathways included calcium and ion channel activity, membrane transport, and depolarization-related processes.

**Figure 4.**
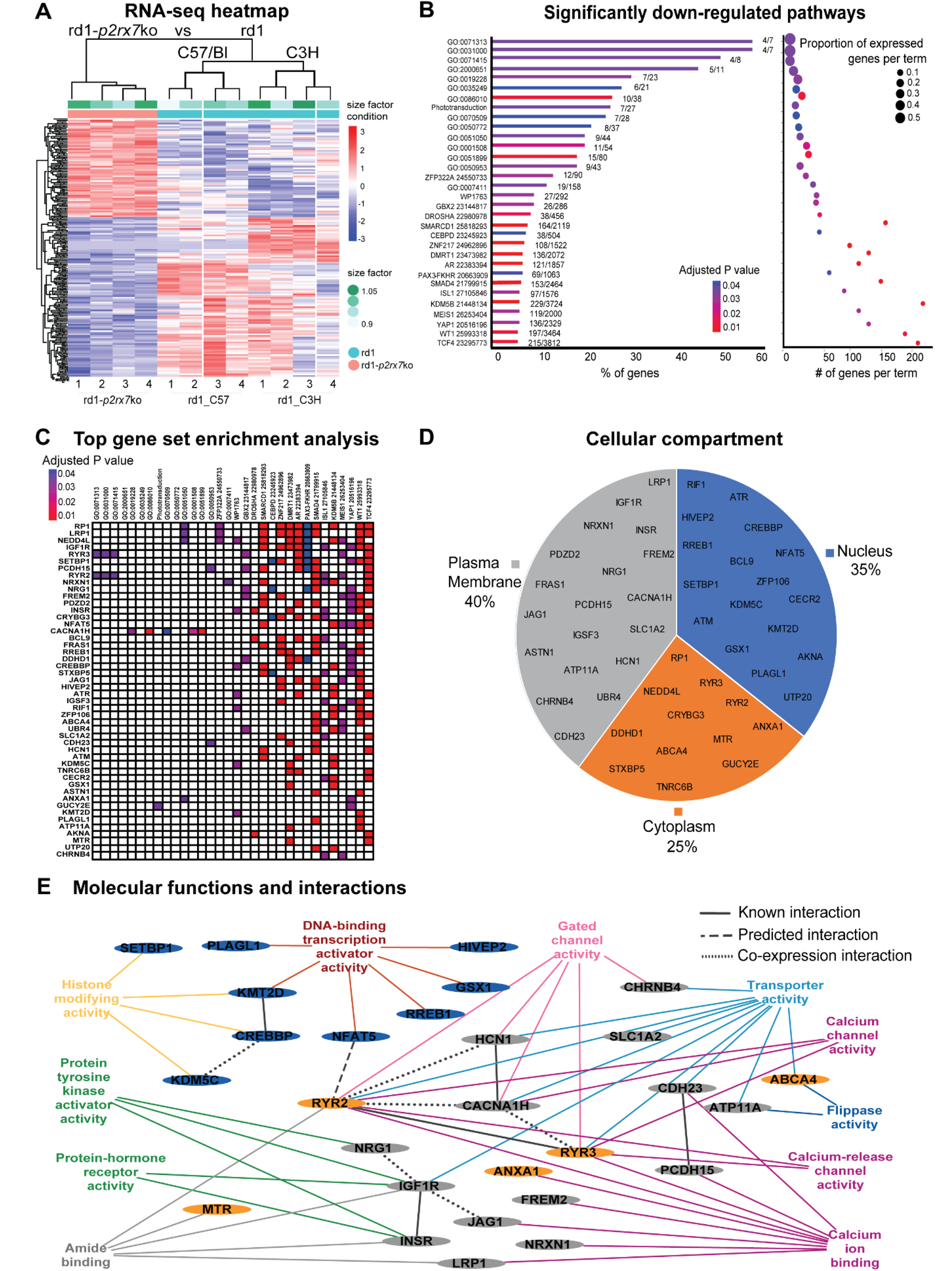
Degeneration-enhanced expression of ion channels is *p2rx7*-dependent. **A)** Heatmap of transcriptional regulation patterns. The image visually represents the differential expression of the top 1000 genes in each retinal sample from rd1-*p2rx7*ko and rd1 mice. Three strains were tested (n=4 per strain), each column represents a sample, each row corresponds to a gene, and each cell displays normalized gene expression values. **B)** KEGG enrichment analysis of significant down-regulated pathways. The figure presents the top 32 significant pathways based on p-adj <0.05 and abs(log2FoldChange) > 0, identified by RNA-seq. The y-axis indicates the pathways identified by their ID (see the readable term name table in supplement materials). The left x-axis demonstrates the percentage of genes, the expressed genes (left number) out of the total genes number (right number) in each GO term. The right x-axis demonstrates the number of genes per term, the bubble size indicates the proportion of expressed genes. The color bar indicates the p- adjusted value, the blue represents a higher value, the red represents lower value. **C)** Gene set enrichment analysis (GSEA) of the top 48 down-regulated genes associated with the 32 significant down-regulated pathways. Pathways are shown horizontally and genes vertically. The colored cells indicate the frequency of gene expression association within each pathway, aligning with its p-adjusted value. **D)** Distribution of top down-expressed genes according to their cellular localization. **E)** Molecular functions and interactions network visualization. The terms and their corresponding connections to the associated genes are colored labeled. Nodes represent genes, blue for nuclear, gray for plasma membrane and orange for cytoplasmic gene localization. The black lines represent the type of protein-protein interaction network analyzed by STRING.

Among the top 50 most significantly and frequently downregulated genes, 48 appeared in two or more GO terms (**Fig. 4C**). Cellular localization analysis revealed that 40% of these genes were plasma membrane-associated (including receptors, transporters, and ion channels), 35% were nuclear (including transcription factors), and 25% were cytoplasmic, including ryanodine receptors and cytoskeletal components (**Fig. 4D**). Functional enrichment using the ToppGene Suite showed that these 48 genes were associated with 275 GO biological process terms, 72 cellular component terms, and 54 molecular function terms. To identify common functional themes, GO terms were ranked by p-value, grouped based on shared genes and functional similarity, and filtered to retain terms containing more than one gene from the top 48. This approach identified 11 major GO term groups interconnected by shared molecular functions (**Fig. 4E**).

Protein–protein interaction analysis revealed physical associations among key downregulated genes, including multiple interactions between ryanodine receptor 2 (RYR2), RYR3, voltage-gated Ca^2+^ channel subunit alpha 1 H (CACNA1H) and hyperpolarization-activated cyclic nucleotide-gated channel 1 (HCN1), which were linked to GO categories such as ion transporter activity, voltage-gated channel activity, and calcium ion binding. These findings are consistent with a role for the RA–P2X7R signaling axis in driving retinal hyperactivity during photoreceptor degeneration. Notably, no significant and consistent changes were observed in the expression of RAR-dependent genes (**Table S1A**) or photoreceptor-associated genes (**Table S1B**) in rd1-*p2rx7*ko retinas as compared to both controls. These results indicate that *p2rx7* deletion does not rescue photoreceptor degeneration nor prevent RAR-dependent transcriptional changes in the surviving inner retina.

### Degeneration-dependent upregulation of *p2rx7* drives increase in HCN1 channel expression and activation in Off-RGCs

We have already shown that transcription of HCN1 is upregulated in the inner retina of rd1 mice and that the ionic current carried by HCN channels is upregulated in rd1 RGCs [13]. To determine if *p2rx7*-dependent signaling is necessary for degeneration-induced changes in HCN expression and activity, we compared the transcriptional status of HCN1-4 (**Fig. 5A-D**) in rd1 and rd1-*p2rx7*ko mouse retinal lysate at P60 and at P120. The expression of HCN1 increased with age and was greatly reduced in retinas from knock-out mice lacking *p2rx7* (**Fig. 5A**). Knocking-out *p2rx7* had a similar, but smaller effect on HCN-2, but only in older mice (p120). In contrast, knocking out *p2rx7* had no effect on HCN3 and HCN4 (**Fig. 5B-D**).

**Figure 5.**
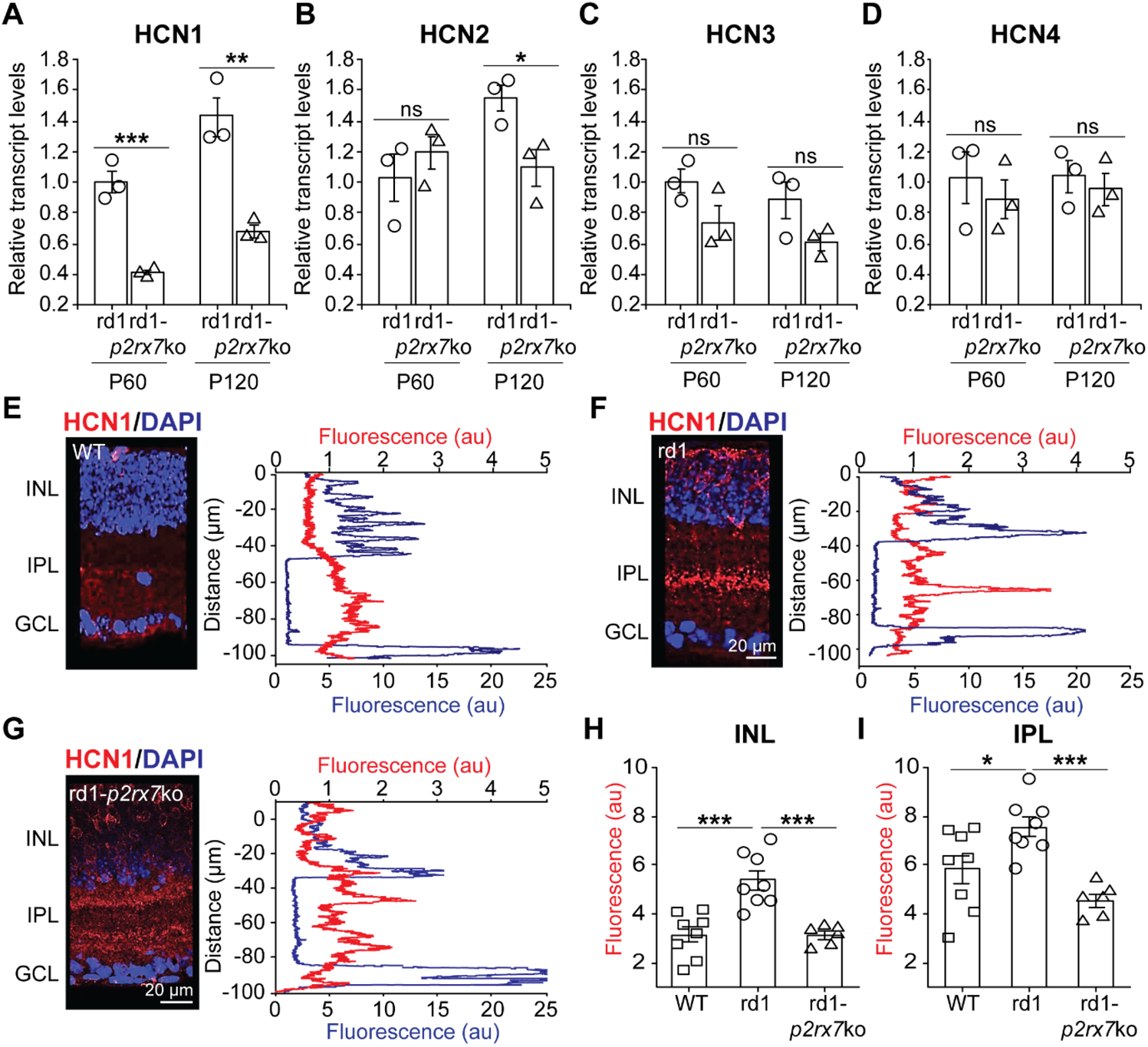
P2RX7-dependent expression of HCN1 in the degenerated retina. **A-D**) Relative transcript levels of HCN1 (**A**), HCN2 (**B**), HCN3 (**C**) and HCN4 (**D**) as assessed by qRT-PCR using the ΔΔCt-quantification method and the housekeeping gene GAPDH, in retinal lysates from 3 rd1 and 3 rd1-*p2rx7*ko mice at P60 and at P120 (total n=12). **E-G**) Immunolabeling of WT (**E**), rd1 (**F**) and rd1-p2×7ko (**G**) retinal slices with a primary antibody against HCN1 revealed with a red-fluorescent secondary antibody and counterstained with the nuclear blue dye DAPI. The staining profile for blue and red fluorescence across the retina, from the outer limit of the INL to the anterior edge of the GCL (∼100 µm wide) is displayed next to each image. **H-I**) Quantification of red fluorescence in the INL (**H**) and IPL (**I**) in the retinas of 8 WT, 8 rd1 and 6 rd1-*p2rx7*ko mice. **A-D, H-I**) Values are shown for individual datapoints (WT – square; circle – rd1; triangle – rd1-p2×7ko), and as mean ± SEM. Wilcoxon test, *p<0.05, **p<0.01, ***p<0.001.

With this information in hand, we next assessed the tissue localization of HCN1 in WT (**Fig. 5E**), rd1 (**Fig. 5F**), and rd1-*p2rx7*ko (**Fig. 5G**) in retinal cryosections. The results reveal that photoreceptor degeneration leads to a *p2rx7*-dependent increase in the expression of HCN1 in both the INL (**Fig. 5H**) and the IPL (**Fig. 5I**), but not in the GCL. Most of the HCN1-immunolabeling appeared in the middle of the IPL, in a tight band at the junction between the *a* and *b* IPL sub-laminae (**Fig. 5F** vs **5G**).

To determine whether increased HCN1 transcription and translation mRNA results in increased HCN-dependent ionic current (I_h_), we obtained whole-cell patch-clamp recordings from RGCs in retinal flat-mounts from WT, rd1 and rd1-*p2rx7*ko mice. Membrane voltage was clamped to -60 mV and cells were subjected to 6 incrementing hyperpolarizing voltage steps (10 mV, 4.0 sec.), changing their membrane potential from -60 mV to -120 (**Fig. 6A**, [24]). Retinal samples were superperfused with saline containing the same mixture of neurotransmitter receptor inhibitors that we used in MEA recordings (**Fig. 3**), and currents elicited by voltage steps were recorded before and after adding ZD7288 (10 µM) an inhibitor of HCN channels (**Fig. 6B** & [25, 26]). The recording pipette was filled with intracellular dye (AF488) to reveal the RGC axon as well as its dendritic stratification within the IPL, using DNA counterstaining (Nuclear ID or DAPI) to determine the position of the somata in the GCL and the INL, and measure the distance between them. If the recorded cell’s dendrites penetrated <50% of the IPL, the cell was classified as a putative On-RGC (**Fig. 6C**), and if >50%, it was classified as a putative Off-RGC (**Fig. 6D**). We omitted bistratified RGCs (‘On/Off’) from this analysis, as well as cells lacking axons.

**Figure 6.**
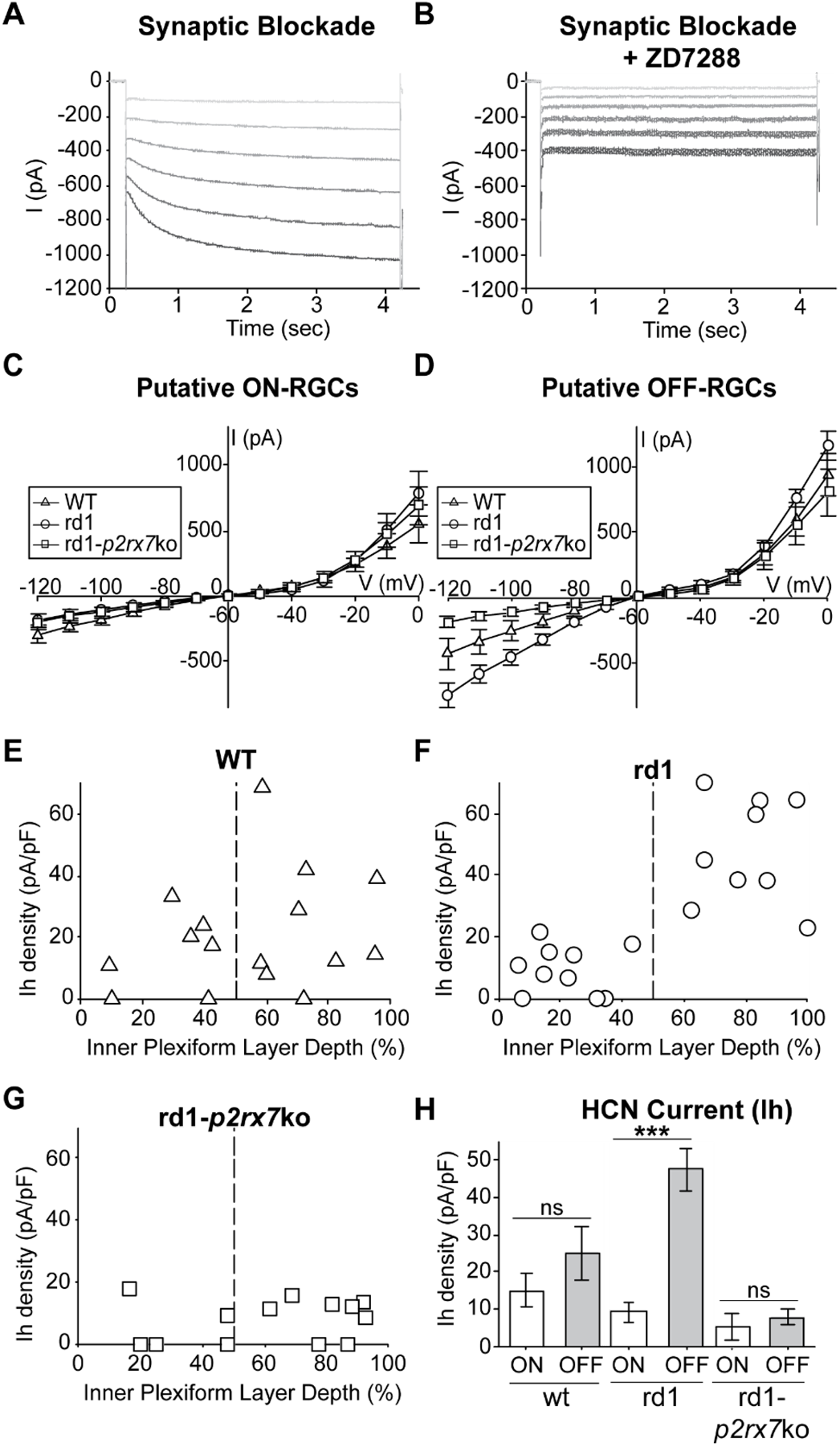
*P2rx7*-dependent increase in HCN currents in the Off-RGCs of degenerated retinas. **A-B**) Current-voltage (IV) curves in putative On- (**A**) and Off-RGCs (**B**), in WT (triangles), rd1 (circles), and rd1-*p2rx7*ko (squares) mouse retinas. **C-E**) Average density of Ih at -110 mV for each recorded RGC in WT (**C**), rd1 (**D**) and rd1-*p2rx7*ko (**E**), plotted against the depth of dendritic penetration in the inner plexiform layer (IPL). **F**) Summary quantification of **C-E**, where Ih density is shown as mean ± SEM in putative On- (white bars) and Off-RGCs (grey bars). Wilcoxon test, ***p<0.001. **A-F**) Recordings from a total of 16 WT-RGCs (On = 7, Off = 9); 19 rd1-RGCs (On = 10, Off = 9); and 13 rd1-*p2rx7*ko-RGCs (On = 5, Off = 8).

The resulting current-voltage (IV) curves show that photoreceptor loss does not significantly alter the electrophysiological responses of On-RGCs, and neither does knocking out *p2rx7*. However, in Off-RGCs, photoreceptor degeneration (rd1) was associated with greatly increased I_h_, but this was absent in rd1-*p2rx7*ko retinas. The surface area of different types of RGCs varies considerably, which could result in different values of I_h_ in different cells. To remove this as a potential confound, we calculated current density (current/capacitance) and plotted this as a function of the dendritic IPL depth, in WT (**Fig. 6E**), rd1 (**Fig. 6F**) and rd1-*p2rx7*ko (**Fig. 6G**) retinas. In the WT retina, absent degeneration, the density of I_h_ did not differ between On- and Off-RGCs. In rd1 retinas, I_h_ density was 2-3 fold larger in Off- than in On-RGCs, or WT-RGCs, regardless of cell identity (**Fig. 6H**). Knocking-out *p2rx7* in rd1 eliminated the increased *Ih* density in rd1 Off-RGCs. These results suggest that RGC hyperactivity in degenerated retinas is generated by Off-RGCs, in which *p2rx7*-dependent expression of HCN1 enhances the I_h_ current, increasing membrane excitability.

### Degeneration-dependent P2X7R-signaling dysregulates RGC Ca^2+^ levels

The results from our RNA-seq experiment (**Fig. 4**) revealed P2X7R-dependent upregulation of HCN1 channels in rd1 retinas (**Fig. 5,6**). The same RNA-seq data also indicated a *p2rx7*-induced increase in the expression of genes involved in Ca^2+^-dependent signaling. Notably, knocking-out *p2rx7* in rd1 mice led to reduced expression of RYR2/3 and CACNA1H (**Fig. 4E**). RYRs mediate release of intracellular Ca^2+^ from the endoplasmic reticulum [27] and they are expressed in many neurons in the inner retina [28, 29]. CACNA1H expression has been demonstrated in type-3 (Off-type) cone bipolar cells (CBCs), contributing to low-threshold depolarization and Ca^2+^ influx, as well as modulation of the amplitude and timing of the b -wave in electroretinogram (ERG) recordings [30–32]. Therefore, it seemed possible that a sustained increase in intracellular Ca^2+^ could link up-regulation of P2X7R to changes in the intrinsic properties of RGCs, leading to hyperexcitability and hyperactivity.

To test whether photoreceptor degeneration alters the intracellular Ca^2+^ concentration in neurons of the inner retina in a *p2rx7*-dependent manner, we measured the fluorescence of the genetically encoded Ca^2+^-indicator GCaMP6f, virally expressed in neurons of the inner retina driven by the human synapsin-1 neuronal promoter (hSyn1) (**Fig 7A**, [33]). Adult WT, rd1 and rd1-*p2rx7*ko mice were intravitreally transduced using AAV2, and retinas were live imaged 1-1.5 months later, in flat-mounted preparations under continuous perfusion of oxygenated physiological solution at 34°C (see **Materials and Methods**). Flat-mounted retinal samples were imaged in 3-4 fields of view, each captured in 3 repetitions of 300 seconds at a 10 Hz (100 msec exposure time per frame), and each containing hundreds of cells (200 x 200 µm). Ca^2+^-dependent fluorescence was imaged under baseline conditions; and again 15 minutes after continuous perfusion with physiological solution supplemented with a mixture of synaptic blockers (**Fig 7B**). We averaged the relative values of Ca^2+^-dependent fluorescence for each cell over the entire time frame. Our results show that *p2rx7* is necessary for degeneration-induced elevated Ca^2+^ in the GCL, independent of synaptic input (**Fig. 7B**), suggesting that functional P2X7Rs, which happen to be Ca^2+^-permeant, are required for heightened Ca^2+^ in GCL neurons.

**Figure 7.**
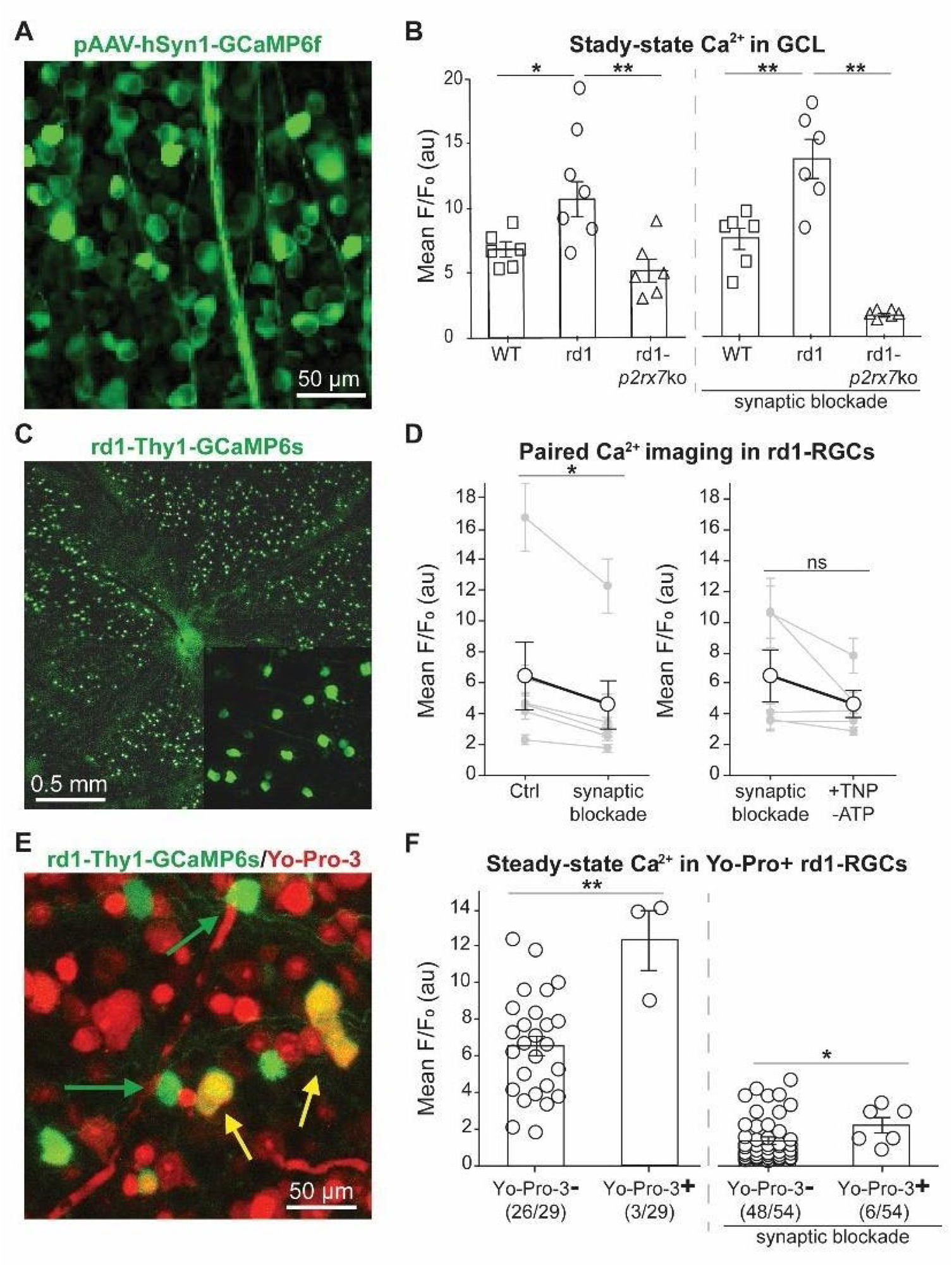
P2X7 receptor signaling sustains elevated steady-state Ca²⁺ levels in rd1 RGCs. **(A)** Confocal image of GCaMP6f fluorescence in the ganglion cell layer (GCL) following intravitreal delivery of pAAV-hSyn1-GCaMP6f. Robust neuronal expression enables measurement of steady-state intracellular Ca²⁺ dynamics in situ. **(B)** Quantification of mean steady-state Ca²⁺ levels (F/F₀, arbitrary units) in WT, rd1, and rd1-*p2rx7*ko retinas under baseline conditions (left) and during synaptic blockade (right). rd1 RGCs exhibit significantly elevated Ca²⁺ compared to WT, which is reduced in rd1-*p2rx7*ko mice. Synaptic blockade does not abolish the Ca²⁺ elevation in rd1 but markedly reduces Ca²⁺ in rd1-*p2rx7*ko. Values are shown as mean ± SEM, *p<0.05, **p<0.01; one-way ANOVA with post hoc Tukey’s HSD; n = 275 - 328 cells from 6 mice and 12 retinas per genotype. **(C)** Wide-field image of rd1-Thy1-GCaMP6s retina illustrating sparse RGC labeling and large-scale imaging field. Inset shows higher magnification of RGCs with axons projecting to the optic disk. **(D)** Paired Ca²⁺ imaging in identified rd1 RGCs before and after pharmacological manipulation. Left, synaptic blockade significantly reduces steady-state Ca²⁺ in a subset of rd1 RGCs. Right, addition of TNP-ATP under synaptic blockade does not further reduce Ca²⁺ levels. Grey lines connect values in individual mice; black circles indicate mean ± SEM. *p<0.05; ns, not significant; paired t-test after Shapiro Wilk normality test; n = 25-40 cells per retina in 10 retinas from 5 mice, per condition. **(E)** Dual imaging of rd1-Thy1-GCaMP6s (green) with Yo-Pro-3 (red) labeling to identify P2X7R-permeable pore formation. Green arrows indicate GCaMP6s⁺/Yo-Pro-3⁻ RGCs; yellow arrows indicate double-positive cells (GCaMP6s⁺/Yo-Pro-3⁺), consistent with P2X7R activation. **(F)** Steady-state Ca²⁺ levels in Yo-Pro-3⁻ and Yo-Pro-3⁺ rd1 RGCs under baseline conditions (left) and during synaptic blockade (right). Yo-Pro-3⁺ RGCs display significantly elevated Ca²⁺ compared to Yo-Pro-3⁻ cells, an effect that persists under synaptic blockade. Values are shown as mean ± SEM with individual recorded cells are shown as circles. *p<0.05, **p<0.01; Mann–Whitney test.

To focus our assessment on RGCs, we crossbred rd1 mice with a transgenic line expressing GCaMP6s exclusively in RGCs [18], as evidenced by the presence of axons projecting to the optic disk in all cells expressing GFP, while also providing sparse distribution of the Ca^2+^-indicator (“rd1-Thy1-GCaMP6s”, **Fig. 7C**). In flat-mount samples, we imaged Ca^2+^-dependent fluorescence in single RGCs before and after blocking all synaptic inputs (**Fig. 7D**, **left panel**), as well as before and after antagonizing all P2X receptors with 100 µM TNP-ATP (**Fig. 7D**, **right panel**). We have already shown that similar conditions (incubation with 100 µM TNP-ATP for 15-30’) prevents Yo-Pro-1 loading in degenerated retinas [13]. These samples, expressing GCaMP6s, were imaged at 2Hz (500 msec exposure time / frame) [33]. Overall, our results show a moderate reduction of ∼25% in Ca^2+^-dependent fluorescence after synaptic blockade or treatment with TNP-ATP, with only one retinal sample showing a significant decrease in Ca^2+^-dependent fluorescence upon addition of TNP-ATP (**Fig. 7D**, **left panel,** *p<0.05). In these experiments, a slow decrease in baseline fluorescence is likely due to gradual photobleaching of GFP throughout the experiment (45-60’ overall after flat-mounting) and prolonged light exposure during tissue imaging. Overall, acute pharmacological blockade of P2X7R does not robustly reduce GCaMP6 signals in the general RGC population, suggesting these receptors contribute little to acute Ca^2+^ influx, and have little immediate excitatory effect on the cells that might cause calcium influx through voltage-gated calcium channels. To explore a potential relationship between P2X7R activation and higher levels of intracellular Ca^2+^ in rd1-RGCs, we repeated these experiments following incubation with Yo-Pro-3, a red-shifted analogue of Yo-Pro-1, to enable Ca^2+^ imaging with GCaMP in hyperpermeable cells, which we have shown are predominantly Off-RGCs (**Fig. 7E**). We found that Yo-Pro-3-positive cells were more likely to have higher Ca^2+^ than Yo-Pro-negative cells (**Fig. 7F, left panel**). This difference was maintained after synaptic blockade (**Fig. 7F, right panel**). Taken together, our results suggest that in response to photoreceptor degeneration, Off-RGCs display increased *p2rx7* expression and P2X7R activation, leading to chronically elevated intracellular Ca^2+^. We propose that over time, elevated Ca^2+^ leads to changes in expression of voltage-gated ion channels including HCN1, contributing to RGC hyperactivity.

## Discussion

Retinal remodeling is a functionally consequential feature of photoreceptor degenerative disorders. Although prior work established that RA-signaling is necessary for the emergence of electrophysiological remodeling, including bipolar cell loss of calcium current and RGC hyperactivity [2,7], the molecular intermediates linking RA-dependent transcription to altered membrane physiology have remained undefined. Here we identify P2X7 receptors as a critical effector in this cascade. We show that degeneration-induced RA signaling transcriptionally upregulates *p2rx7*, that P2X7R expression is necessary and sufficient for RGC membrane hyperpermeability and hyperactivity, and that genetic deletion of *p2rx7* prevents both acute and chronic remodeling phenotypes. Together, these findings position P2X7R as a mechanistic bridge between RA-dependent transcriptional reprogramming and maladaptive plasticity in the retina.

### RA signaling directly engages the purinergic stress axis

The purinergic signaling pathway is a form of cell–cell communication in which purine nucleotides activate two types of receptors for ATP/ADP: the ionotropic P2X channels and the metabotropic P2Y G-protein coupled receptors [34]. The nucleoside adenosine, a breakdown product of ATP, also activates P1, another group of G-protein coupled receptors. Together, activation of these receptors regulates intracellular Ca²⁺, membrane potential, and second-messenger pathways, thereby influencing neurotransmission, synaptic plasticity, inflammation, and cell survival in the nervous system. Dysregulated purinergic signaling, elicited by cellular stress such as nutrient or oxygen deprivation, contributes to neuronal and glial retinal pathophysiology through mechanisms such as excessive Ca²⁺ influx, neuroinflammation, and microglial activation. Over time, these processes can lead to neurodegeneration [35–37]. In the retina, ATP released by neurons, Müller glia, and retinal pigment epithelium activates purinergic receptors in multiple retinal cell types [38, 39]; overactivation of receptors such as P2X7R can drive Ca²⁺-dependent neuronal death, gliosis, and degeneration in diseases including glaucoma, diabetic retinopathy, retinitis pigmentosa, and age-related macular degeneration [37, 40, 41].

Our data support a direct transcriptional link between RA signaling and *p2rx7* expression. The presence of RAR-binding motifs in the *p2rx7* promoter, coupled with the rapid reduction in P2X7R protein levels following intraocular RAR blockade, strongly suggests that RA/RAR induces *p2rx7* transcription rather than signaling indirectly through secondary stress pathways, such as excessive Ca²⁺ influx and neuroinflammation. The tight correlation between RAR-reporter activity and Yo-Pro-1 uptake further reinforces the coupling between RA-dependent transcription and functional P2X7R macropore formation.

P2X7R has long been conceptualized as a sensor of extracellular ATP and metabolic distress. In immune and glial cells, its activation triggers ion conduction, plasma membrane depolarization, Ca²⁺ influx, inflammasome activation, and downstream transcriptional cascades [14, 15]. In retinal disease models—including glaucoma, diabetic retinopathy, and inherited dystrophies—P2X7R expression and activation are elevated and associated with inflammation and neuronal vulnerability [16, 42–46]. However, these models have largely framed P2X7R as responding to heightened extracellular ATP.

Our findings revise this framework. In rd1 retinas, acute increase in ATP is not sufficient to explain RGC hyperactivity; rather upregulation of P2X7R expression is also required [7, 13]. RA signaling is both necessary and sufficient to drive the upregulation. Thus, rather than functioning solely as a passive ATP sensor, P2X7R in the degenerating retina is transcriptionally engaged by RA signaling and thereby primed to amplify purinergic tone. This shifts P2X7R from a downstream effector of metabolic stress to an active participant in remodeling initiated by photoreceptor loss.

### P2X7R role in acute permeability and chronic hyperactivity

A central insight of this study is the observation that acute pharmacological blockade and genetic deletion of *p2rx7* have distinct effects. Acute antagonism of P2X receptors suppressed Yo-Pro-1 permeability but failed to reduce RGC hyperactivity. In contrast, genomic deletion abolished both. This distinction suggests that P2X7R contributes to acute membrane hyperpermeability via macropore opening, consistent with its canonical ionotropic function; while RGC hyperactivity is due to P2X7R driving long-term transcriptional reprogramming, resulting in coordinated changes in RYR2/3, CACNA1H, and HCN1, along with other voltage-gated channel subunits, pointing toward potential reconfiguration of intrinsic cell excitability and Ca^2+^ dynamics.

Calcium influx through excitatory ligand-gated ion channels, including neuro- and gliotransmitter receptors, initiates signaling cascades that can further amplify intracellular calcium dynamics and promote long-term synaptic changes. For example, activation of NMDA receptor channels allows substantial Ca²⁺ entry into postsynaptic neurons, leading to activation and autophosphorylation of the Ca²⁺ sensor and effector CaMKII, which enhances synaptic efficacy through phosphorylation and recruitment of Ca²⁺-permeant AMPA glutamate receptors, thereby creating a positive feedback loop that reinforces excitatory transmission and long-term potentiation (LTP) [47–50]. Metabotropic glutamate receptors, especially mGluR1 and mGluR5, also elevate intracellular Ca²⁺ indirectly through phospholipase C activation and release of Ca²⁺ from intracellular stores in the endoplasmic reticulum, triggering changes in transcriptional regulation and inducing long-term depression (LTD) of synaptic transmission [51–53]. Calcium-permeable nicotinic ACh receptors channels, particularly α7-containing isoforms, can similarly couple neuronal depolarization to downstream transcriptional pathways [54–56]. Collectively, these Ca²⁺-dependent signaling mechanisms converge on nuclear effectors including CREB and immediate early genes such as cFos, linking transient synaptic activity to persistent changes in gene expression, neuronal adaptation, and structural plasticity [57, 58].

Similarly, P2X receptors regulate neuronal and glial gene expression through Ca²⁺-dependent signaling pathways that couple extracellular ATP release to transcriptional remodeling during both physiological activity and neurodegeneration. Activation of P2X receptors Ca²⁺-influx and downstream engagement of MAPK, calcineurin, and PKC signaling cascades, leading to activation of transcription factors including CREB, NFAT, NF-κB, AP-1 family members (c-Fos/c-Jun), and Egr proteins in neurons and glia, thereby linking purinergic signaling to inflammatory, stress-response, and plasticity-associated transcriptional programs [59–61]. Although P2X7Rs are most abundantly expressed in microglia and other immune-associated glial populations, neuronal P2X7R expression has been reported in the hippocampus, cortex, retina, dorsal root ganglia, and presynaptic terminals, where receptor density is generally lower under physiological conditions but markedly increases during inflammation, injury, excitotoxicity, and retinal degeneration [62, 63]. Moreover, we have shown P2X7R up-regulation in RGCs in photoreceptor-degenerated mice [13]. Neuronal P2X7Rs can localize to somata, dendrites, axon terminals, and peri-synaptic regions, where they can modulate neurotransmitter release, intracellular Ca²⁺-dynamics, and excitability in a chronic manner. While acute activity-dependent gene expression induced by transient Ca²⁺-influx through excitatory neurotransmitter receptors like NMDA and AMPA activates immediate-early genes such as c-Fos, Arc, and Egr1 within minutes; sustained P2X7R activation during chronic extracellular ATP accumulation produces prolonged Ca²⁺-dysregulation, reactive oxygen species generation, inflammasome activation, and persistent NF-κB/MAPK signaling in neurons and glia, driving long-term inflammatory transcriptional states, gliosis, cytokine production, and degeneration-associated gene expression programs characteristic of chronic CNS and retinal disease [64]. Our study highlights the role of P2X7Rs as chronic participants in a signaling pathway that brings about long-term changes, underlying maladaptive plasticity to cellular stress and tissue degeneration.

### Remodeling without rescue of photoreceptors

The P2X receptor family comprises seven isoforms (P2X1–P2X7) that assemble as homo- or hetero-trimeric receptors with distinct biophysical and signaling properties, of which all but P2X1R, have been reported to express in the retina, specifically in photoreceptors, Müller glia, microglia, retinal pigment epithelium, and RGCs [65–69]. Although P2X7R is the principal isoform associated with inflammatory signaling, Ca²⁺ overload, and degeneration, other retinal P2X receptors may also contribute to synaptic transmission, glio-transmission, mechanosensation, and ATP-dependent stress responses through heteromeric assembly and modulation of Ca²⁺ permeability [65, 70, 71]. Among these receptors, P2X7 is considered unique because prolonged or repeated activation by high extracellular ATP concentrations can induce formation of a large non-selective membrane pore (“macropore”) permeable to molecules up to ∼900 Da [13, 72], promoting ionic dysregulation, inflammasome activation, cytokine release, mitochondrial dysfunction, and cell death. Yet, some evidence also suggests that P2X2 and P2X4 receptors may also exhibit a macropore under sustained stimulation [73–75].

In this study, the genomic deletion of *p2rx7* in rd1 mice affected with a severe and aggressive form of RP did not prevent photoreceptor degeneration, only the downstream changes associated with inner retinal remodeling. Therefore, in the rd1 mouse, degeneration and remodeling can be mechanistically separable processes. This distinction is critical for therapeutic design: preventing photoreceptor loss could be much more difficult than repressing remodeling, and even if both strategies were possible, they could be complementary rather than conflicting interventions. However, it is important to notice that the changes to the inner retina may occur on a spectrum of possibilities: from homeostatic and compensatory rewiring when a small fraction of cones are artificially ablated [3–5]; to visually-detrimental pathophysiological remodeling triggered by RA when a significant portion of rods degenerate [2, 7, 22, 76], to exacerbated loss of RGCs in aged and blind mice [77]. Our results suggest that RA–P2X7R signaling is a key inflection point in the transition from minimal compensatory rewiring to “full-blown” pathological remodeling. By linking photoreceptor loss to transcriptional reprogramming of RGC firing patterns, this pathway may define when adaptive neuronal plasticity becomes maladaptive.

### Therapeutic implications for vision preservation and restoration

Current gene and cell therapies for RP aim to preserve or repopulate photoreceptors, but they are ineffective once degeneration has progressed beyond a critical stage. Vision restoration strategies—including stem cell transplantation, optogenetics, azobenzene photoswitches, and electronic implants —rely on a functional inner retina capable of encoding and transmitting meaningful signals to the brain [78–80]. RGC hyperactivity reduces the signal-to-noise ratio, substantially degrading retinal circuit computations [2, 7].

Our data suggest that targeting P2X7R or its downstream transcriptional effectors may be a valid approach to preserve circuit fidelity during disease progression. Inhibiting RA synthesis improves light response signal-to-noise ratios in partially degenerated retinas [2, 7], and P2X7R modulation may extend this benefit by preventing both membrane hyperpermeability and long-term activity changes. However, the differential effects of acute versus genetic manipulation indicate that durable intervention—potentially via gene therapy, sustained pharmacological inhibition, or transcriptional modulation—will be required to meaningfully suppress remodeling if targeting P2X7R.

Beyond RP, P2X7R has been implicated in epilepsy, neuroinflammation, and Alzheimer’s disease, conditions also characterized by maladaptive plasticity [14, 15]. Our findings raise the possibility that P2X7R-mediated transcriptional reprogramming represents a widespread mechanism coupling extracellular stress signals to intrinsic neuronal remodeling. Comparative studies across CNS regions may reveal whether RA signaling similarly engages purinergic pathways in other degenerative contexts.

### A model for RA–P2X7R–driven remodeling

We propose the following model: photoreceptor degeneration elevates RA signaling [2, 7], leading to RAR-dependent transcriptional upregulation of *p2rx7*. Increased P2X7R expression enhances membrane permeability and Ca²⁺ influx, initiating both acute ionic changes and chronic transcriptional cascades. These cascades reconfigure ion channel expression (e.g.: HCN1) and Ca²⁺ handling machinery, stabilizing a hyperactive RGC phenotype. In this framework, P2X7R is not simply a biomarker of stress but a necessary intermediate that converts RA-dependent transcription into pathophysiological remodeling. By identifying this link, our study clarifies how photoreceptor loss is translated into circuit-level dysfunction and provides a defined molecular entry point for therapeutic intervention.

## Acknowledgements

This work was completed using funds available to R.H.K (NIH grants P30EY003176 and R01EY024334) and to M.T. (NIH grants P30EY08098, R01EY036020-01A1 and Research to Prevent Blindness Career Development Award), as well as Research to Prevent Blindness unrestricted gift, and the generous support from the Eye and Ear Foundation, to the Dept. of Ophthalmology, University of Pittsburgh.

## Author contributions

M.T. and L.A. designed, performed, and analyzed the results for most experiments and wrote the manuscript. J. E. supported immunohistochemistry experiments and mouse genotyping. B. D. supported MEA experiments. R.H.K. designed the study, wrote and edited the manuscript.

## STAR METHODS

### RESOURCE AVAILABILITY

#### Lead contact

Further information and requests for resources and reagents should be directed to the lead contact, Dr. Richard H Kramer.

#### Materials and availability

This study generated a unique RAR-reporter construct which will be made available to researchers upon request to Dr. Michael Telias.

#### Data and code availability

- Sequencing data are available at GEO (ID: GSE325143).
- This paper did not generate any original code.
- Any additional information required to reanalyze the data reported in this paper is available from the lead contact upon request.

### EXPERIMENTAL MODEL AND SUBJECT DETAILS

#### Animals

All animal procedures were approved by the UC Berkeley Institutional Animal Care and Use Committee (IACUC) and by the University of Rochester Committee on Animal Resources (UCAR) and conformed to NIH guidelines.

Wild-type (WT) mice (C57BL/6J; Jackson Laboratory, JAX:000664; RRID:IMSR_JAX:000664) and rd1 mice (C3H/HeJ; JAX:000659; RRID:IMSR_JAX:000659) were obtained from The Jackson Laboratory. P2rx7 knockout mice (B6.129P2-P2rx7^tm1Gab^/J; JAX:005576; RRID:IMSR_JAX:005576) were crossed with rd1 mice to generate rd1-P2rx7^−/−^ mice (‘rd1-*p2rx7*ko’) on a mixed background. rd1-Thy1-GCaMP6s mice were generated by crossing rd1 mice with C57BL/6J-Tg(Thy1-GCaMP6s)GP4.3Dkim/J (JAX:024275; RRID:IMSR_JAX:024275).

Genotyping was performed according to Jackson Laboratory protocols using genomic DNA extracted from ear or toe biopsies and AccuPower PCR Pre-Mix (Bioneer). Unless otherwise specified, mice were 1–4 months of age (P30–P120). Both male and female mice were used. Animals were housed under a 12-hour light/dark cycle with food and water provided ad libitum (≤5 mice per cage).

## METHOD DETAILS

### Intravitreal Injections

Mice were anesthetized with 2% isoflurane. Pupils were dilated using 1% tropicamide and 2.5% phenylephrine. Topical anesthesia was achieved with 0.5% proparacaine. A scleral incision posterior to the ora serrata was made using a 30G needle, and 1–2 μL of solution was injected intravitreally using a blunt 33G Hamilton syringe. Tobramycin (0.3%) was applied post-procedure.

To inhibit retinoic acid receptor (RAR) signaling, BMS-493 was injected at a final intraocular concentration of 0.1–0.5 μM. Vehicle consisted of PBS containing 1% DMSO.

For calcium imaging experiments, AAV2 vectors encoding hSyn1-GCaMP6f (≥10^13^ gc/mL) were injected at P30–P45.

### Retinal Dissection

Eyes were enucleated immediately following euthanasia. Retinas were isolated in oxygenated artificial cerebrospinal fluid (ACSF) containing (in mM): 119 NaCl, 2.5 KCl, 1 KH₂PO₄, 1.3 MgCl₂, 2.5 CaCl₂, 26.2 NaHCO₃, and 20 D-glucose, equilibrated with 95% O₂/5% CO₂.

For live imaging and electrophysiology, retinas were flat-mounted on nitrocellulose filter paper. For molecular analyses, retinas were transferred immediately to lysis buffer. For immunohistochemistry, eyes were fixed in 4% paraformaldehyde (PFA).

### RNA Sequencing

Total RNA was extracted from retinas of rd1-p2rx7ko (n=4) and rd1 mice (n=8; 4 C3H background, 4 C57 background; P60–P90) using the RNeasy Plus Mini Kit (Qiagen, Cat#74134). RNA quality was assessed spectrophotometrically (DeNovix DS-11+).

Libraries were prepared from 200 ng RNA using the TruSeq Stranded mRNA Library Preparation Kit (Illumina, Cat#20020595) and sequenced on a NovaSeq 6000 platform.

Reads were demultiplexed using bcl2fastq (v2.20.0) and processed using FastP (v0.23.1). Differential expression analysis was performed in R (v3.5.1) using DESeq2 (v1.34.0). Adjusted p values (Benjamini–Hochberg) < 0.05 were considered significant. Log₂ fold changes were moderated using lfcShrink.

Only genes consistently regulated across both rd1 background comparisons were retained for downstream analysis. Functional enrichment was performed using EnrichR and ToppGene. Protein interaction networks were generated using STRING and visualized in Cytoscape.

### RT-qPCR

Both retinas from each mouse were pooled. RNA was extracted (RNeasy, QIAGEN) and reverse transcribed (500 ng RNA; SuperScript III, Thermo Fisher). Quantitative PCR was performed using SYBR Green Master Mix (Applied Biosystems) on a BioRad thermocycler.

Relative expression was calculated using the ΔΔCt method with Actb as housekeeping gene. Primer sequences are listed below and in Table S1.

### Western Blot

Protein was extracted from individual retinas using RIPA buffer (Thermo Fisher). Western blotting was performed by Boster Bio (Pleasanton, CA) in a blinded manner. Membranes were probed for P2X7R and GAPDH.Band intensities were quantified using FIJI (ImageJ) and normalized to GAPDH.

### Immunofluorescence

Eyes were fixed in 4% PFA (1.5 h, RT), cryoprotected in 10–30% sucrose, embedded in OCT, and cryosectioned at 14 μm thickness. Sections were blocked in PBS containing 2.5% BSA and 0.1% Triton X-100. Primary antibodies were applied overnight at 4°C; secondary antibodies were applied for 1 h at room temperature. Sections were mounted with DAPI-Fluoromount G and imaged using a Nikon laser-scanning confocal microscope.

### Yo-Pro Uptake Assay

Flat-mounted retinas were incubated in oxygenated ACSF (34°C) with 200 nM Yo-Pro-1 (15 min) and Nuclear-ID Red (1:500; 3 min). Z-stacks (1.5 μm step size) were acquired spanning the GCL through the inner nuclear layer.

### Calcium Imaging

Retinas expressing hSyn1-GCaMP6f or Thy1-GCaMP6s were imaged at P60–P90 using a spinning-disk confocal microscope (Olympus BX51WI). Excitation was 470–490 nm; emission was collected at 515–550 nm. Images were acquired at 1–5 Hz using a 40× water-immersion objective. Fluorescence was quantified as ΔF/F₀ = (F − F₀)/F₀.

### Multielectrode Array Recordings

Retinas were placed ganglion-cell-layer-down on a 60-electrode MEA (MEA 1060-2-BC, Multi-Channel Systems). Signals were filtered at 200 Hz and digitized at 20 kHz. Spikes exceeding 4 SD above baseline were detected. Spike sorting was performed using principal component analysis (Offline Sorter, Plexon).

### Patch-Clamp Recordings

Cell-attached recordings were obtained using a MultiClamp 700B amplifier at −60 mV under continuous ACSF perfusion (34°C). Data were acquired using pCLAMP 10.4 and analyzed with ClampFit 10.5.

### Pharmacology

Synaptic blocker cocktail (μM): 10 AP4, 40 DNQX, 30 AP5, 10 SR-95531, 50 TPMPA, 10 strychnine, 50 tubocurarine.

P2X receptor antagonist: 100 μM TNP-ATP.

HCN channel blocker: 10 μM ZD7288.

## QUANTIFICATION AND STATISTICAL ANALYSIS

Data are presented as mean ± SEM. Statistical analyses were performed using GraphPad Prism. Normality was assessed using Shapiro–Wilk tests. Outliers were identified using the Thompson Tau method. Parametric comparisons were performed using Student’s t tests or ANOVA with Tukey HSD post hoc testing. Nonparametric comparisons were performed using Wilcoxon rank-sum tests. Bootstrapping was used when group sizes were unequal. Statistical significance was defined as p<0.05.

## KEY RESOURCES TABLE

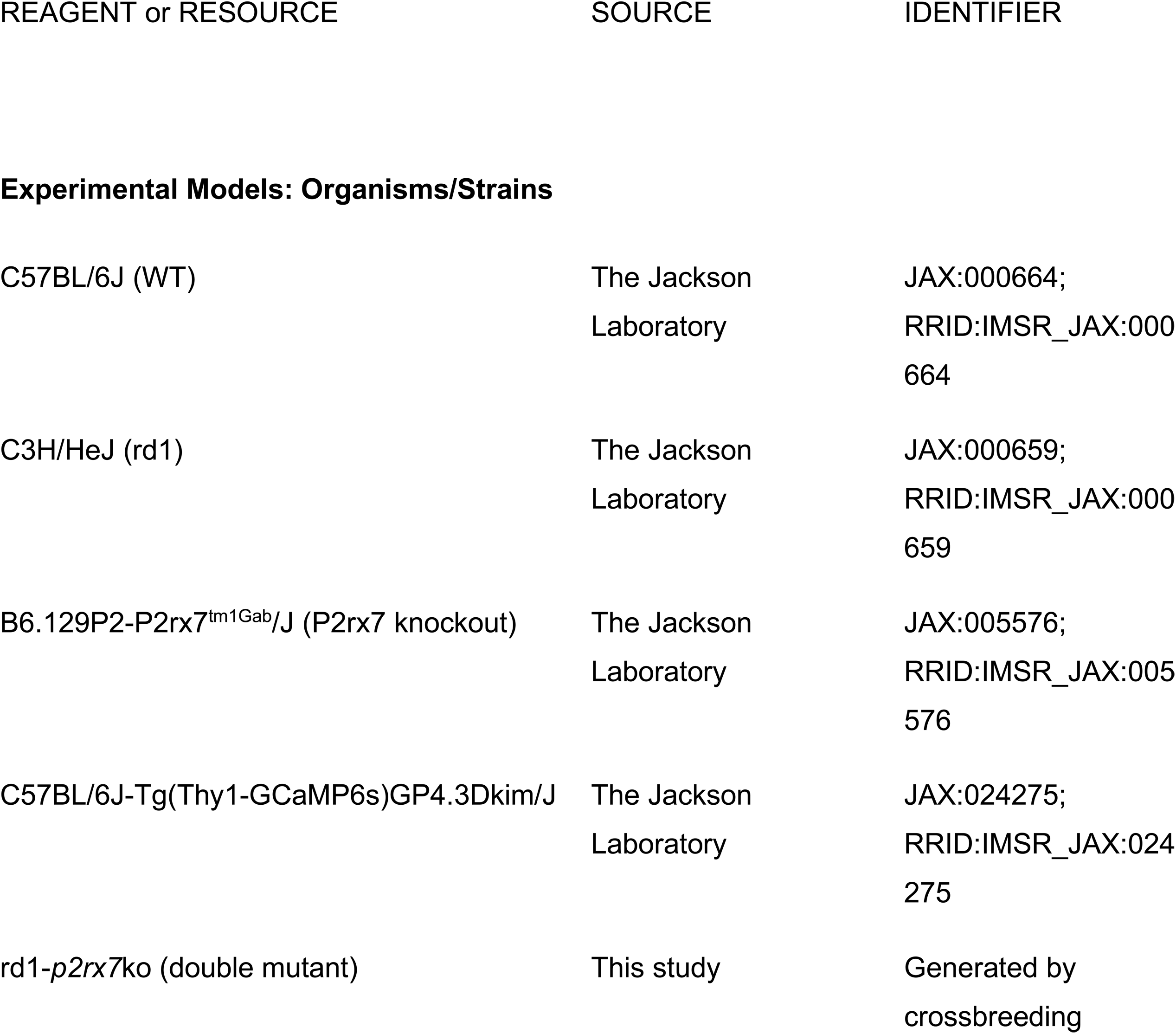
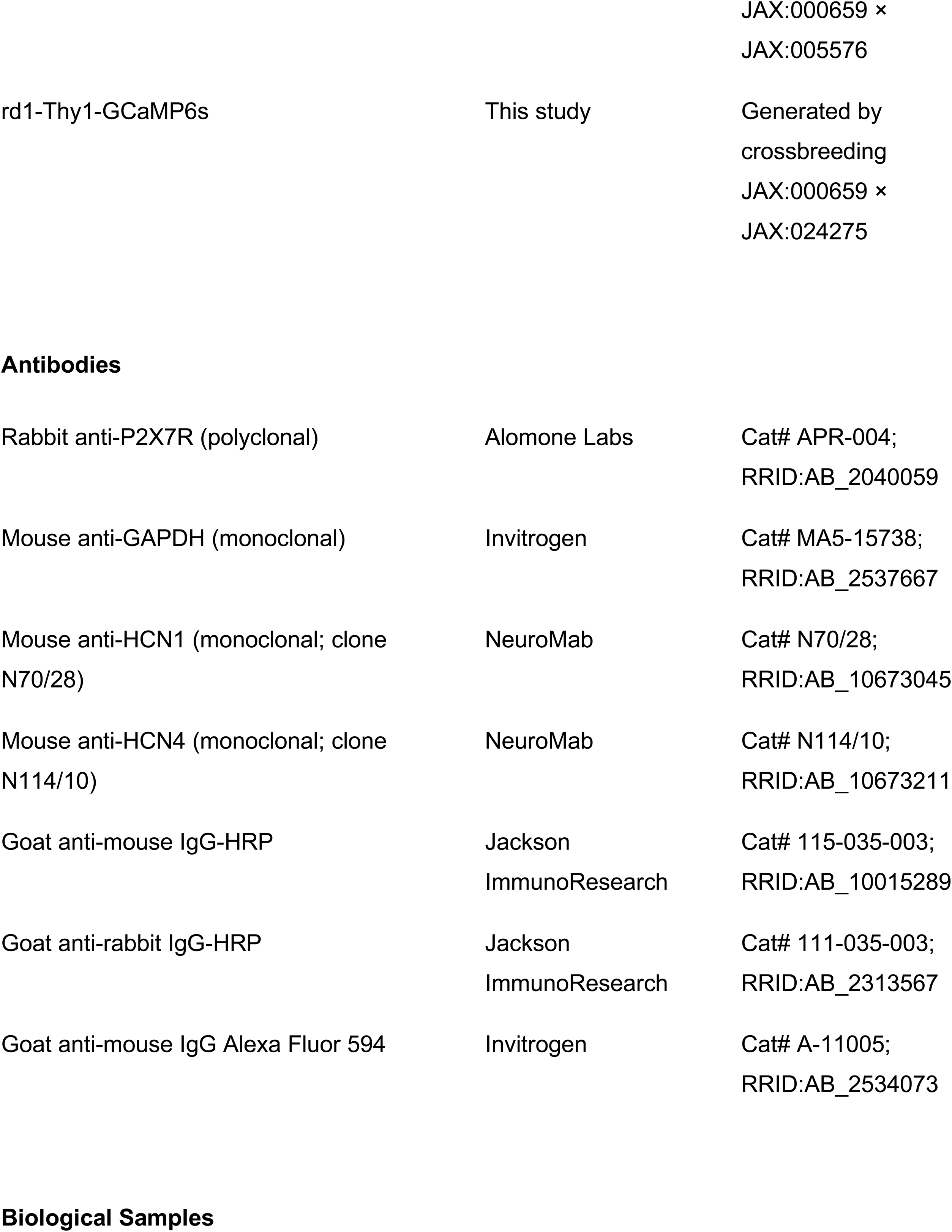
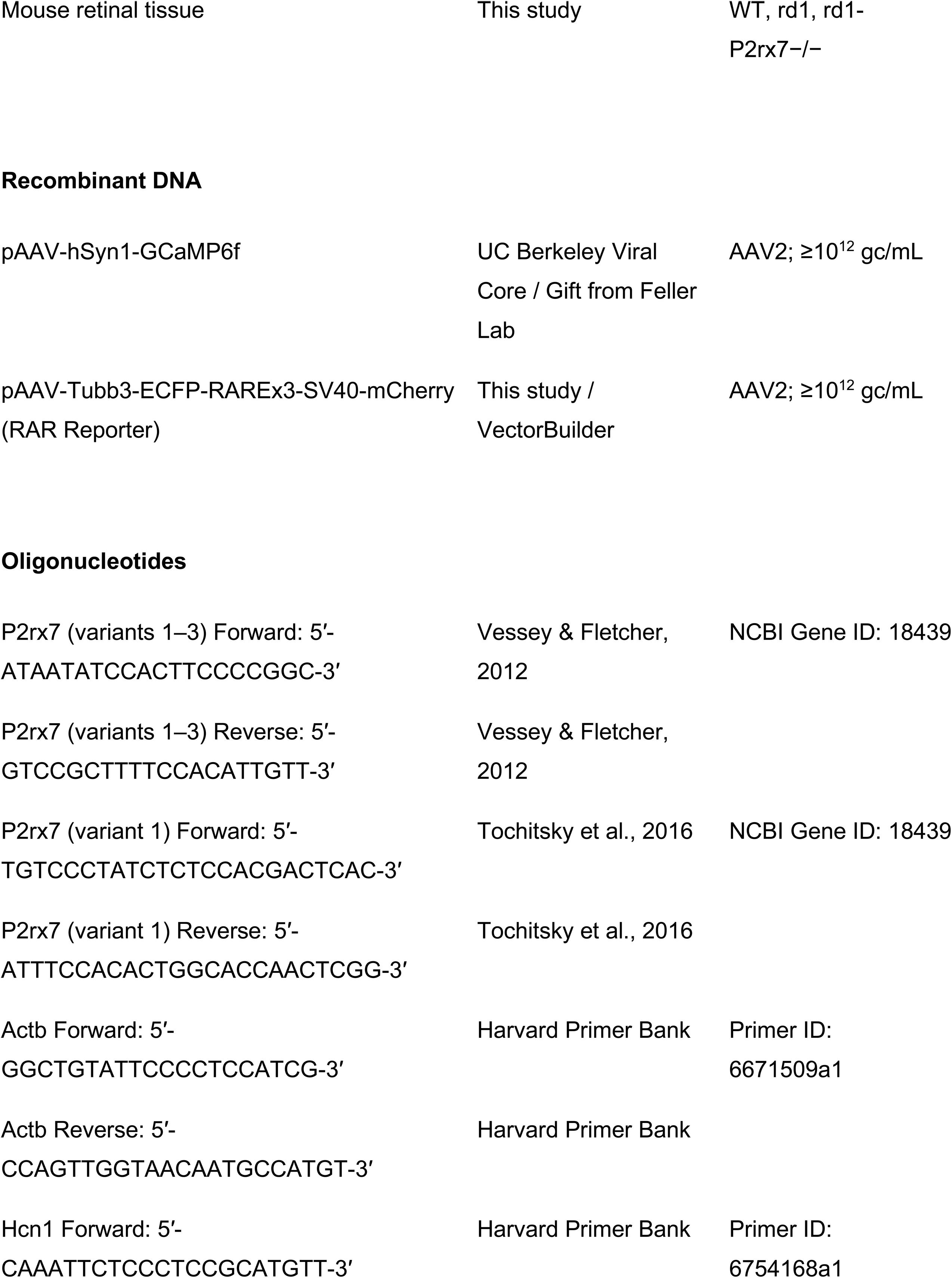
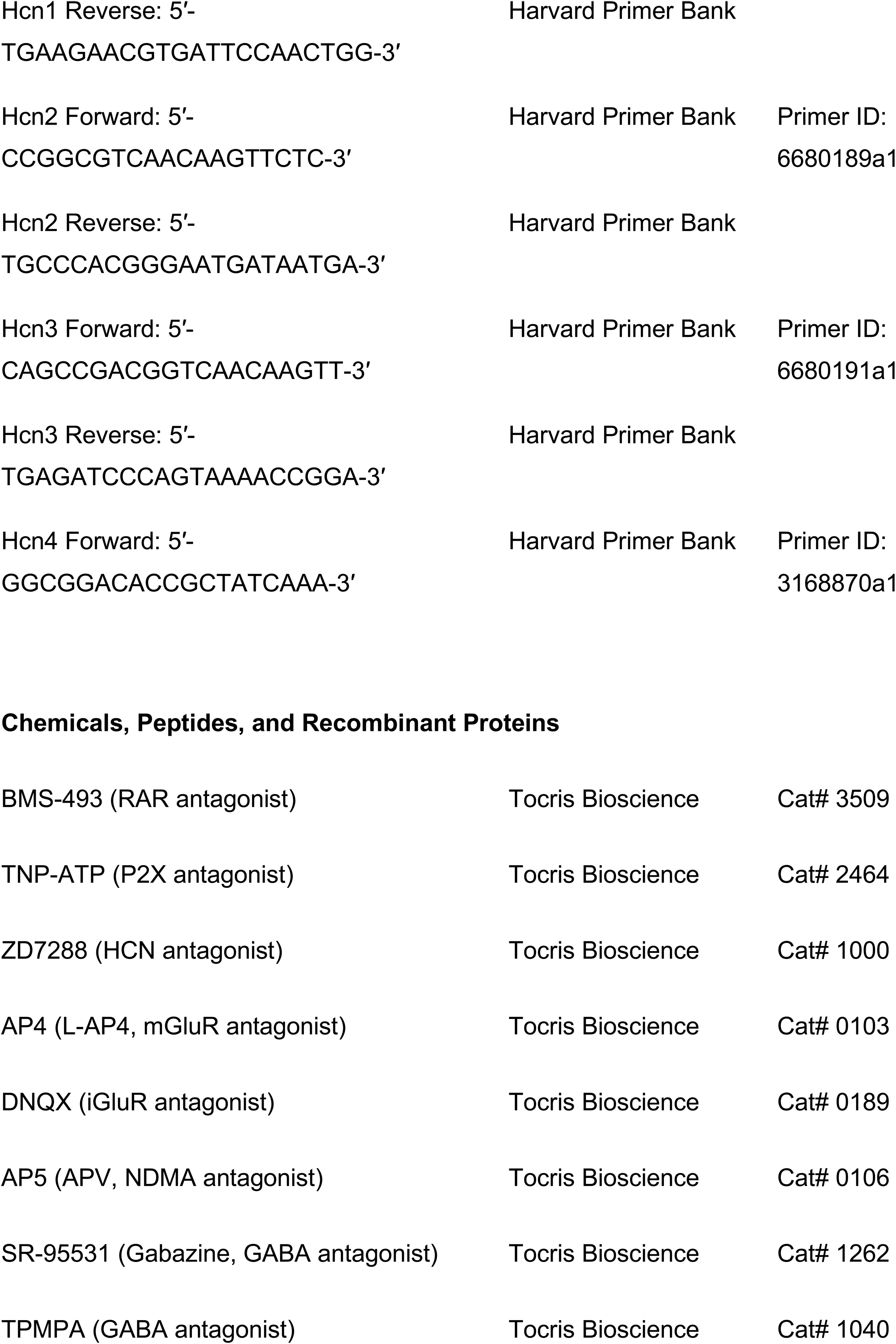
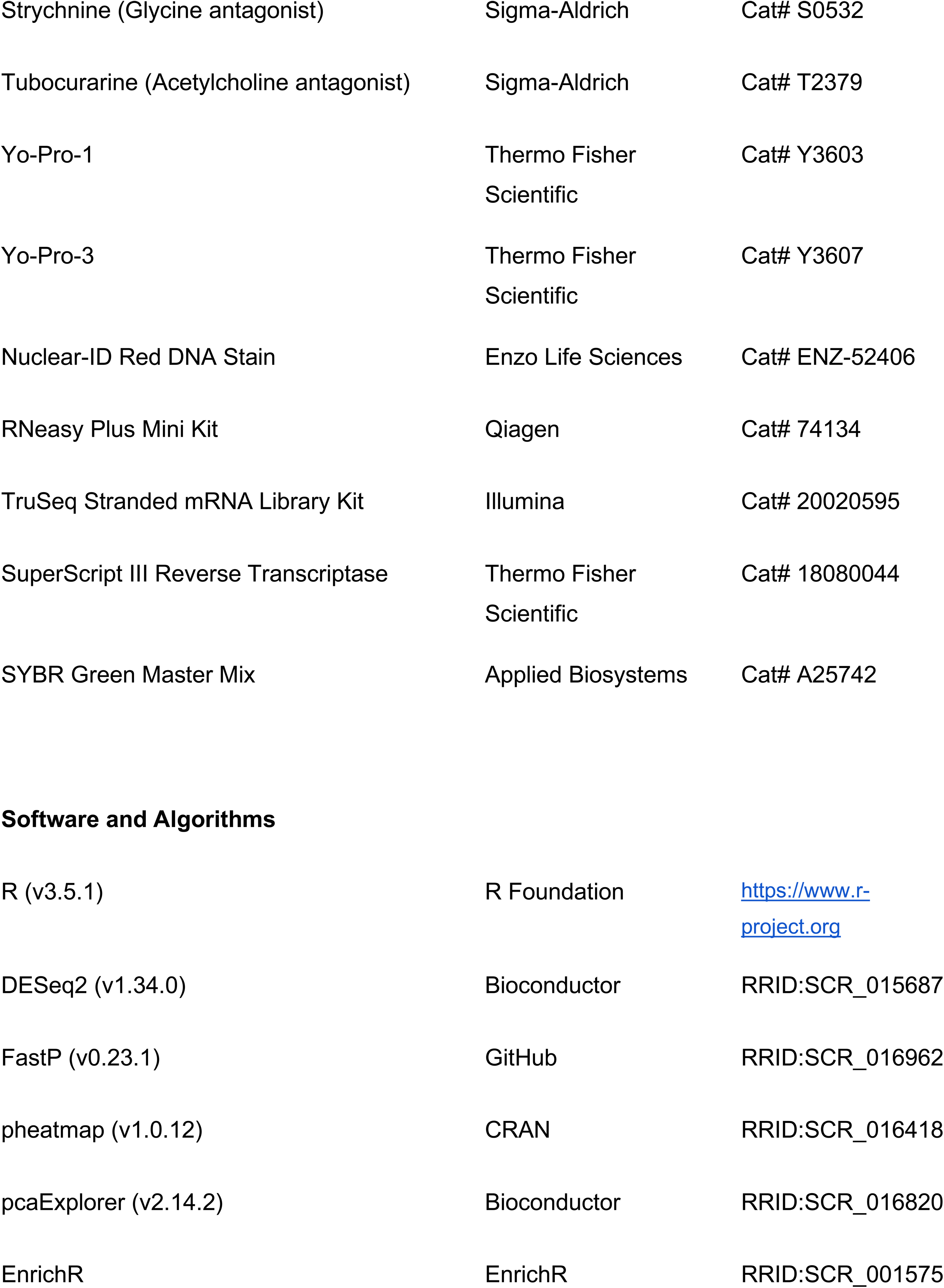
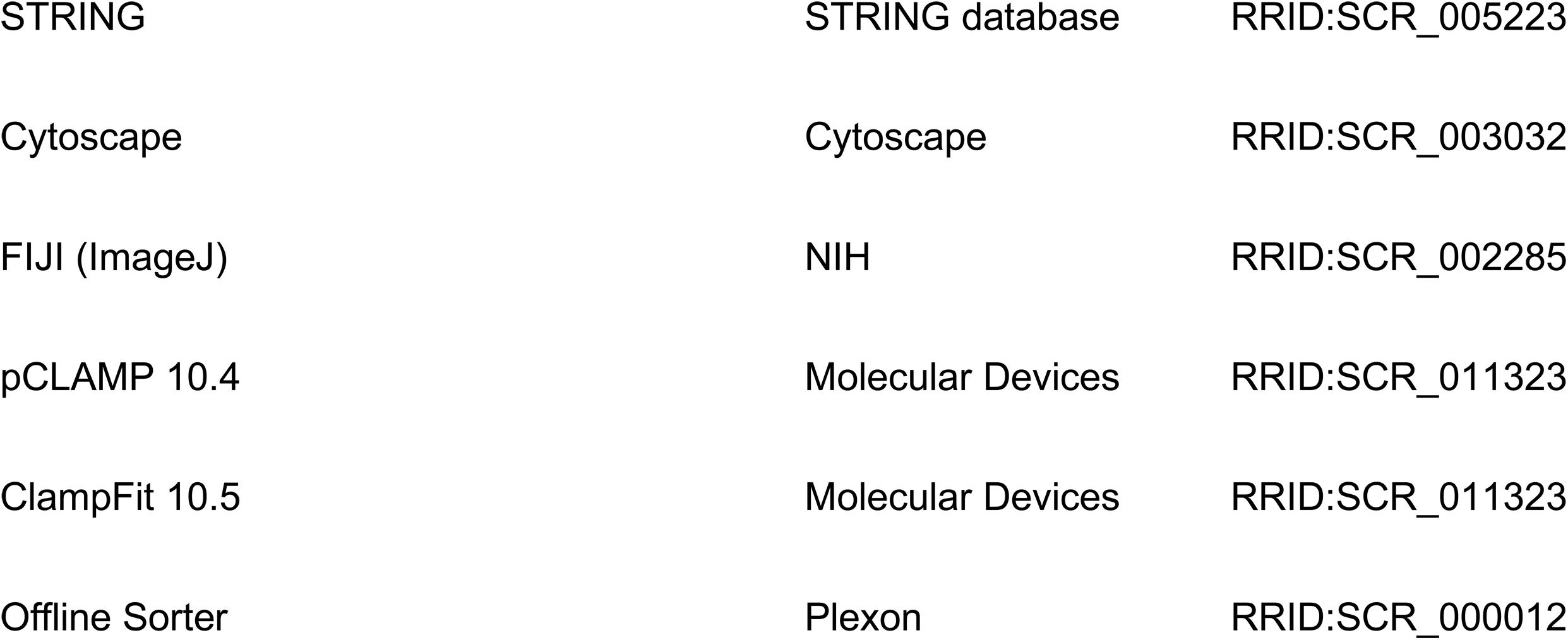

