## Supplementary figures and images for "The P2X7 receptor is an intermediate in the retinoic acid-signaling pathway that induces neuronal remodeling in retinitis pigmentosa"

### Supp Figure 1

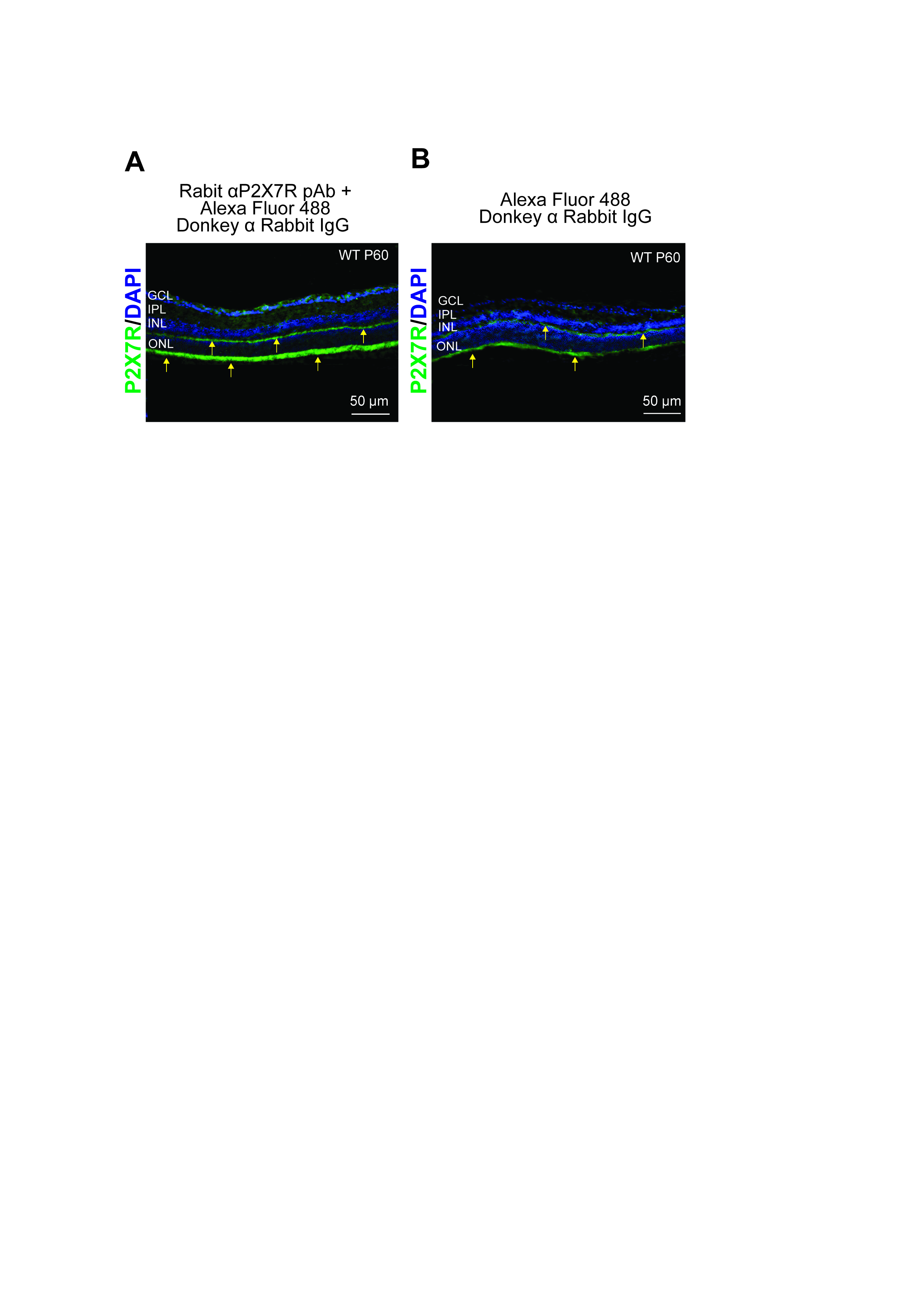

### Supp Figure 2

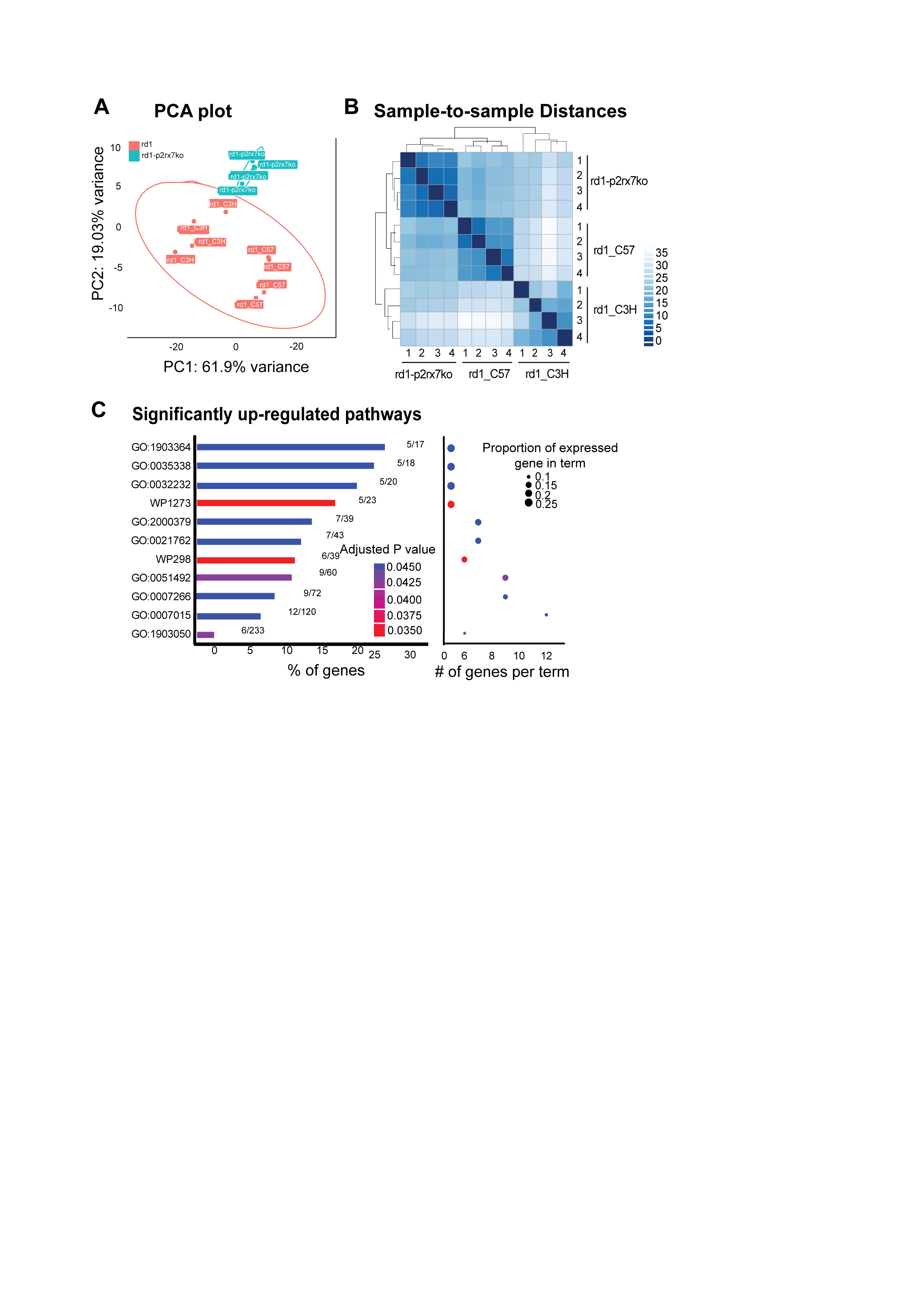
